# A comprehensive analysis of calreticulin mutants reveals distinct biophysicochemical proprieties with a potential for refined targeted therapies

**DOI:** 10.64898/2026.06.19.733337

**Authors:** Orhan Nedim Kurt, Mustafa Baran Ozturk, Eylul Civelek, Ilyas Chachoua

**Affiliations:** Department of Molecular Biology and Genetics, Bilkent University, Ankara, Türkiye

**Author notes:** Equal contribution (Mustafa Baran Ozturk, Eylul Civelek). The authors declare no potential conflicts of interest.

**Keywords:** Calreticulin mutants, Type 1, Type 2, biophysicochemical properties, MPN, MPL

## Abstract

Calreticulin mutations in myeloproliferative neoplasms result in the replacement of the C-terminus acidic sequence with a positively charged tail that causes pathological activation of the thrombopoietin. The two canonical variants are Type-1 and Type-2. The remaining are mainly classified as Type-1 or Type-2 like based on the wild type sequence retained. Here, we performed in silico biophysicochemical analyses of 76 CALR exon 9 frameshift variants by their sequence and predicted biophysical properties, complemented by structural modeling of the mutant homodimers. Beyond confirming the Type-1 versus Type-2 distinction, we found that the Type 1-like variants form a continuum of charge architecture along which two reproducible subgroups can be identified, rather than sharply separated classes. This work refines the conventional mechanism-based classification into a charge-resolved framework and provides testable hypotheses linking novel-tail chemistry to receptor activation in CALR-mutant neoplasms and paves the way for improved targeted therapies based on individual mutants’ characteristics

## Introduction

Myeloproliferative neoplasms (MPNs) are clonal hematopoietic stem cell disorders characterized by aberrant expansion of one or more myeloid lineages (1). Among the driver mutations underlying these malignancies, somatic frameshift mutations in exon 9 of CALR, encoding calreticulin, represent the second most frequent molecular event after JAK2 V617F, occurring in approximately 20–25% (ET) and 25–30% (MF) of all patients (2–4). In its physiological state, calreticulin is a crucial, calcium-binding lectin chaperone that ensures proper folding of glycoproteins and maintains calcium homeostasis within the ER lumen (5). However, pathogenic exon 9 frameshifts universally displace the native CALR C-terminal KDEL endoplasmic reticulum retention signal and replace it with a novel positively charged C-terminal sequence, which aberrantly activates the thrombopoietin receptor (TpoR/MPL) through a mechanism requiring homodimerization of the mutant protein driving consecutive JAK-STAT signaling and subsequent oncogenic transformation

The variants are conventionally classified into two types. Type 1, exemplified by a 52 bp deletion (p.L367fs46), is associated with myelofibrotic transformation and produces a highly cationic novel C-terminal tail, whilst being strongly associated with a high-risk primary or post-ET myelofibrotic transformation, clinically marked by rapid bone marrow remodeling and cytopenias. Conversely, type 2, exemplified by a 5 bp insertion (p.K385fs47), is associated with a more inert proliferative essential thrombocythemia (ET) phenotype characterized by extreme thrombocytosis but a significantly lower risk of fibrotic progression or thrombotic events and generates a distinct tail bearing acidic clusters (6–8). This sequence and clinical diversity implies that the physicochemical properties of the novel C-terminal tail vary substantially across the variant landscape, yet systematic characterization of this variation has remained limited (9).

This clear link between surface charge and disease severity suggests that the standard Type 1/Type 2 binary overlooks a more nuanced continuum of receptor activation. Because current prognostic models lump all non-canonical “Type 1-like” variants into a single risk category, critical differences in patient outcomes remain hidden. By breaking down this Type 1-like cohort by their precise biochemical features, we can evaluate whether subtle shifts in charge distribution push a patient toward a more mild or aggressive prognosis.

Prior computational and biochemical studies have established that the novel CALR C-terminus functions as an intrinsically disordered region (IDR) whose charge patterning, rather than a defined three-dimensional structure, is central to TpoR engagement (10). The canonical Type 1 versus Type 2 binary captures the most prominent axis of variation but may obscure biologically meaningful heterogeneity within each class, particularly within the Type 1-like group, which encompasses a broader range of frameshift positions and correspondingly diverse novel tail sequences. Whether this within-class sequence diversity translates into distinct physicochemical and predicted structural properties has not been systematically examined across a curated multi-variant dataset. Here we present a purely computational characterization of 76 CALR exon 9 frameshift variants. Using a comprehensive sequence feature pipeline comprising 101 features across eight biochemical categories, including charge architecture, intrinsically disordered protein (IDP) biophysical metrics, position-specific charge patterning, and kinase consensus motif analysis, we describe three sequence-defined variant groups: Type 2 (n=32), and two exploratory subgroups within the Type 1-like group, designated Type 1-A (n=25) and Type 1-B (n=19). These subgroups are recovered by unsupervised clustering of sequence features. ColabFold v1.5 homodimer predictions are concordant with this subdivision in the rank-1 models; however, multi-model analysis (five models per variant) shows that the predicted compact/extended geometry is reproducible at the Type-1-versus-Type-2 level but is conformationally labile within the Type-1 subgroups. The Type 1-A/Type 1-B distinction is therefore sequence-defined rather than independently validated by structure. Our findings provide an exploratory, sequence-based framework for CALR variant subclassification that extends the canonical two-type model and generates hypotheses for future experimental and clinical investigation. The Type 1-A/Type 1-B subdivision is sequence-defined and should be regarded as hypothesis-generating pending experimental validation.

## Materials and Methods

### Variant dataset

Wild-type calreticulin (CALR) protein sequence was retrieved from UniProt (accession P27797). Variant data for CALR mutations were obtained from CALR-ETdb (dsimb.inserm.fr/CALR-ET). Mutant protein sequences were retrieved directly from the database’s precomputed FASTA files. All variants were screened for the presence of the conserved (to be called anchor) 22-amino-acid sequence (RRMMRTKMRMRRMRRTRRKMRR) in their novel C-terminal tail, shared across majority of pathogenic CALR frameshift mutations. Only anchor-confirmed 76 frameshift variants were retained for downstream analysis: 44 Type 1-like variants associated with myelofibrosis (MF) and 32 Type 2-like variants associated with essential thrombocythemia (ET). The anchor sequence served as the structural reference point for all subsequent position-specific analyses (full sequences in Supplementary Table S1).

### Sequence feature computation

A total of 101 features were computed for each variant organized into eight categories: (A) novel tail amino acid composition, charge metrics (NCPR; net charge per residue, FCR fraction of charged residues), sequence entropy, and tail length; (B) charge patterning via SCD (sequence charge decoration and kappa from localCIDER v0.1.21 (11,12); (C) propensity scales including mean Kyte-Doolittle hydropathy (13), Chou-Fasman beta-sheet propensity (14), and the charge–hydropathy (CH) disorder-boundary metric (15); (D) position-specific formal charges and Grantham distances (16), (E) cysteine geometry; (F) sequence motifs, (G) post-KKRK region metrics for Type 2 variants; and (H) wild-type C-domain fragment length and charge contrast. The complete per-variant feature matrix is provided in Supplementary Table S2, and full definitions of all features, scales, and computational procedures in Supplementary Table S4.

### Unsupervised clustering

Unsupervised clustering was applied exclusively to the 44 Type 1-like variants to identify potential subgroups within this class. (Type 2 variants showed a single, homogeneous cluster, Supplementary Figure S2). The Sγ–Sγ (Gamma sulfur-Gamma Sulfur) inter-chain distance derived from homodimer structural predictions (see below) was designated as a post-hoc validation target and was explicitly excluded from all clustering inputs. A total of 87 features were retained for clustering. terms. All features were standardized to zero mean and unit variance using StandardScaler prior to clustering.

Cluster structure was assessed using the silhouette coefficient across k=2 to k=7 and the gap statistic (17) for k=1 to k=3 (50 reference datasets) (Supplementary Figure S1A). Two clustering algorithms were applied in parallel: k-means (k=2; n_init=50; random_state=42) and agglomerative hierarchical clustering with Ward linkage (17) and Euclidean distance (Supplementary Figure S1D). The two resulting subgroups were designated Type 1-A (n=25) and Type 1-B (n=19).

As a sensitivity analysis, clustering was repeated after (a) excluding all 12 approach zone features requiring imputation and (b) removing one feature from each of 11 near-perfectly correlated pairs. A third analysis applying aggressive decorrelation recovered the partition with ARI = 0.662, reflecting the removal of genuinely informative features rather than mere computational redundancy. For each sample the two-cluster k-means partition was recomputed and matched to the original clusters by maximal Jaccardcy.

Post-hoc feature importance was assessed by Mann-Whitney U test (two-sided) with Benjamini-Hochberg (BH) false discovery rate correction applied across all 83 testable features (4 of 87 features were excluded from testing due to insufficient non-missing values in one subgroup).

Cohen’s d was computed as a descriptive effect size measure. Principal component analysis (PCA) was performed on the standardized feature matrix (Supplementary Figures S1E-F). Cluster stability was further assessed by nonparametric bootstrap (1,000 resamples; per-cluster Jaccard similarity f). For each sample the two-cluster k-means partition was recomputed and matched to the original clusters by maximal Jaccard overlap.

### Sequence-based conformational ensemble prediction

To estimate conformational dimensions independently of structure prediction, we applied the ALBATROSS deep-learning model (via the sparrow package) to the novel C-terminal tail of each variant. ALBATROSS predicts the radius of gyration (Rg), end-to-end distance, and apparent scaling exponent (ν) of disordered sequences from sequence alone.

### Structural predictions

All structural predictions were generated using ColabFold v1.5 (18), AlphaFold2 neural network architecture (19) with MMseqs2-based multiple sequence alignment. For each variant, the unrelaxed rank-1 model was selected for all downstream structural metric extraction, as unrelaxed ColabFold outputs preserve the raw neural network-predicted geometry prior to force-field minimization.

### Structural metric extraction

Structural metrics were extracted from raw PDB. Four monomer metrics were extracted per variant: mean pLDDT across all residues, mean pLDDT over the novel tail region, mean pLDDT over the wild-type region (residues upstream of the frameshift position), and the predicted template modelling score (pTM). 12 homodimer metrics were extracted per variant, comprising Sγ–Sγ and Cb-Cb inter-chain distances and chi1 dihedral angles at the conserved sequence+32 cysteine, inter-chain Ca-Ca contacts within the novel tail (8.0 Angstrom cutoff, normalized by tail length), homodimer novel tail pLDDT, PAE-based confidence metrics (mean, inter-chain, and intra-chain PAE), dimer-pTM, and ipTM. Per-variant structural metrics are provided in Supplementary Table S3, and full metric definitions in Supplementary Table S5.

### Structural group validation

Post-hoc validation of sequence-derived clusters against structural geometry was performed using the Sγ–Sγ inter-chain distance Variants were classified into structural geometry groups based on Sγ–Sγ distance: compact (Sγ–Sγ < 20 Angstrom; n=26) and extended (Sγ–Sγ > 50 Angstrom; n=17), with one variant occupying an intermediate position Correspondence between sequence-derived cluster membership and structural geometry group was assessed by Fisher’s exact test and the adjusted Rand index (ARI).

### Statistical analysis

Group comparisons were performed using the two-sided Mann-Whitney U test. Standardized effect sizes were computed as Hedges’ g (bias-corrected) with pooled standard deviation, reported alongside the nonparametric test statistic. For features with near-zero within-group variance in one subgroup, standardized effect sizes inflate to extreme values (|g| > 10) and are reported as variance-driven artifacts rather than interpreted as effect magnitudes. Three-group comparisons were assessed by Kruskal-Wallis test (20). Multiple testing correction used the Benjamini-Hochberg procedure applied independently within each comparison family.

## Data and Code Availability

All analysis code is openly available at https://github.com/onedimkurt/calr-frameshift-analysis-public, comprising the complete computational pipeline; sequence-feature computation, unsupervised clustering, ColabFold structural-metric extraction, and review-stage validation analyses, together with the variant sequence inputs required to reproduce the sequence-derived and clustering results. The ColabFold structure predictions (monomer and homodimer PDB models, predicted aligned error matrices, and per-residue confidence and score files for all 76 variants) are archived at Zenodo (https://doi.org/10.5281/zenodo.20752925). Full reproduction instructions are provided in the repository README and docs/PIPELINE.md.

## Software and resources

Analyses were performed in Python 3.10.14 (RRID:SCR_008394) using NumPy 1.26.4 (RRID:SCR_008633), SciPy 1.13.1 (RRID:SCR_008058), pandas 2.2.2 (RRID:SCR_018214), scikit-learn 1.5.1 (RRID:SCR_002577), Biopython 1.87 (RRID:SCR_007173), Matplotlib (RRID:SCR_008624), and localCIDER 0.1.21 (Holehouse et al., 2017; idptools/localcider). Sequence-ensemble dimensions were estimated with ALBATROSS via sparrow (Lotthammer et al., 2024; idptools/sparrow). Structure predictions used ColabFold v1.5 (RRID:SCR_025453) with AlphaFold2 (RRID:SCR_025454) and MMseqs2 (RRID:SCR_008184); the AlphaFold Protein Structure Database (RRID:SCR_023662) was used as indicated. Variant annotations were drawn from COSMIC (RRID:SCR_002260), UniProt (RRID:SCR_002380), and the CALR-ETdb database.

## Results

### Dataset composition and universal sequence features

To characterize the biophysicochemical properties of the mutant CALR C-terminal sequences, we computed a panel of 101 sequence-derived features for all 76 variants across eight categories; novel-tail composition and charge metrics (NCPR, FCR), charge patterning (SCD and κ via localCIDER v0.1.21, propensity and physicochemical scales (Kyte–Doolittle hydropathy, Chou–Fasman β-sheet propensity, the charge–hydropathy disorder-boundary metric, and Grantham distance), position-specific formal charges, cysteine-geometry indicators, sequence motifs, and wild-type C-domain features (Methods; Supplementary Tables S2, S4). Applied across the full variant set, this analysis first revealed a series of invariant sequence landmarks shared by all 76 variants.

A cysteine residue at anchor-relative position +32 and +36 was present in all 76 variants, confirming the invariance of these structural markers across the full variant landscape irrespective of primary class (Figure 1A). Because the anchor region and entire post-anchor region are sequence-identical across all variants, all sequence diversity among variants resides exclusively in the approach zone, the segment immediately N-terminal to the anchor.

**Figure 1.**
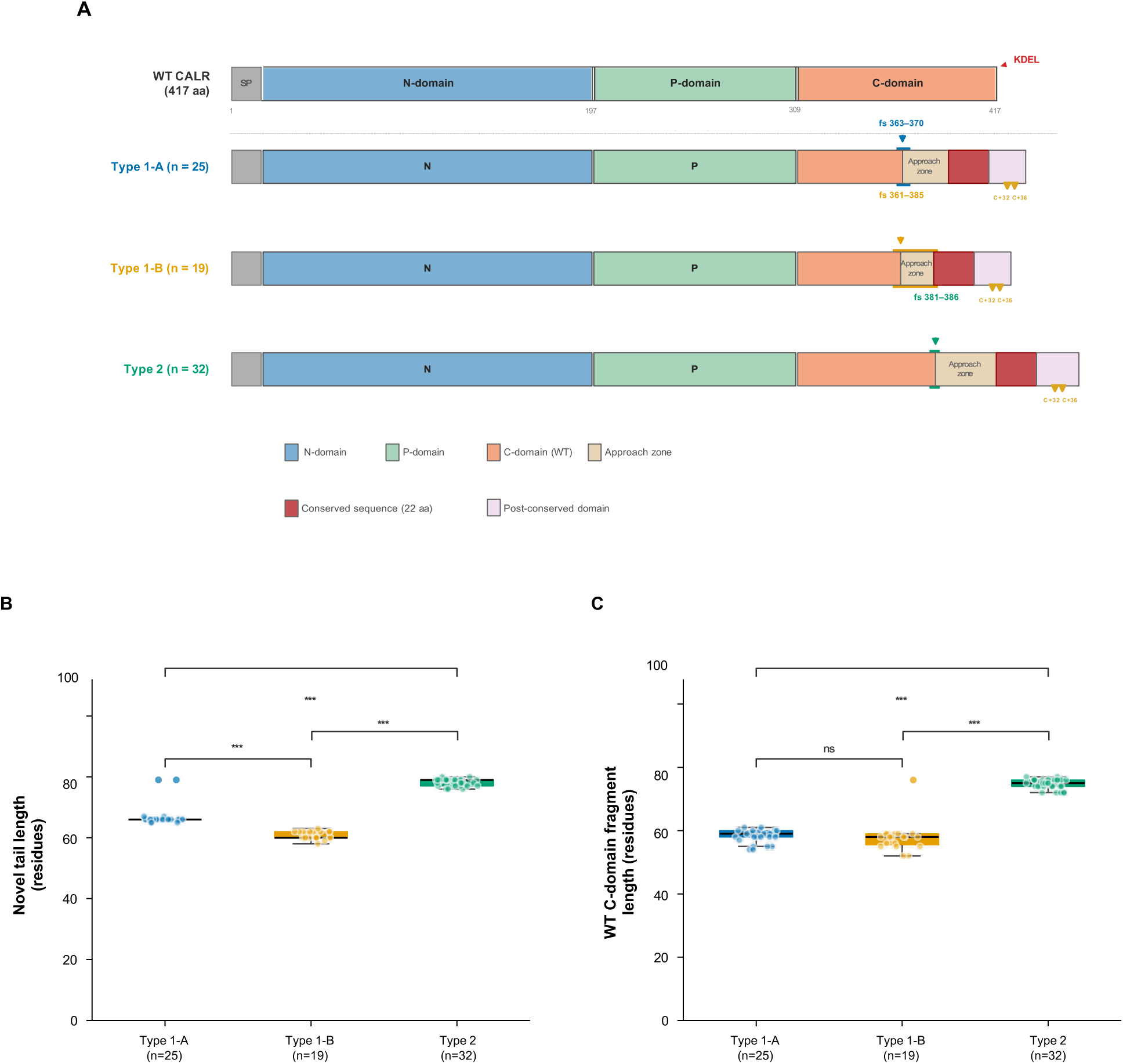
Domain architecture and structural length metrics of 76 CALR exon 9 frameshift variants. (A) Schematic representation of wild-type calreticulin (WT CALR, 417 aa) and the three computationally defined variant groups. The WT protein comprises a signal peptide (SP), N-domain (residues 1–197), P-domain (197–309), and C-domain (309–417) terminating in the ER-retention signal KDEL. Each variant group shares an identical conserved anchor sequence (22 aa, dark red) and invariant cysteines at anchor-relative positions +32 and +36 within the post-conserved domain (lavender). Variant-specific sequence diversity is confined to the approach zone (tan), the segment immediately N-terminal to the anchor. Frameshift positions are indicated by colored arrowheads: Type 1-A (fs 363–370, blue), Type 1-B (fs 361–385, orange), Type 2 (fs 381–386, teal). (B) Novel tail length (residues) across the three variant groups. Type 2 variants have the longest novel tails (mean 78.19 ± 1.15, range 76–80), followed by Type 1-A (mean 67.00 ± 3.64, range 65–79) and Type 1-B (mean 60.84 ± 1.34, range 58–63); Kruskal-Wallis H = 62.35, p = 2.90 × 10⁻¹⁴, ε² = 0.827; all pairwise comparisons ***p < 0.001. (C) Wild-type C-domain fragment length (residues from C-domain start to frameshift position). Type 2 retains substantially more WT C-domain (mean 74.81 ± 1.53) than Type 1-A (58.16 ± 2.08) and Type 1-B (57.89 ± 4.92); the Type 1-A versus Type 1-B difference is non-significant, confirming that both Type 1 subgroups share an essentially identical conserved scaffold. Box plots: median and IQR; whiskers 1.5× IQR; individual points shown. ***p < 0.001; ns, not significant (BH-corrected Mann-Whitney U).

The KKRK tetrapeptide motif was present in all 32 Type 2-like variants and absent from all 44 Type 1-like variants, constituting a perfect discriminator between primary classes at the sequence level. In Type 2-like variants, the KKRK motif was invariably followed by a strongly anionic post-KKRK segment immediately downstream of the sequence previously suggested to be a nuclear localization signal in TET3, CUL4B and Rac1 proteins (21–23). Type 2-like variants also carried exactly three consecutive Asp/Glu clusters of length >=3 in all 32 cases, compared to Type 1-like variants (range 0-1). Type 2-like variants further exhibited an invariant consecutive anionic run of exactly 5 residues in all 32 cases, compared to a mean of 2.87 in Type 1-like variants, and carried exactly two KEE motifs in all 32 cases, absent from virtually all Type 1-like variants. Together with the invariant KKRK motif and three consecutive acidic clusters, these findings define a highly constrained Type 2 sequence architecture.

Type 2-like variants retained a substantially longer wild-type C-domain fragment upstream of the frameshift position compared to Type 1-like variants (Figure 1C), preserving a larger portion of the native CALR acidic C-domain with potential implications for calcium-binding and chaperone function.

### Unsupervised clustering identifies two sequence-defined subgroups within Type 1-like variants

Unsupervised clustering of the 44 Type 1-like variants on 87 standardized features yielded a two-cluster partition that was reproducible across algorithms (k-means and Ward, 100% agreement). Bootstrap resampling indicated that the membership boundary between the two subgroups is soft rather than sharp: per-cluster Jaccard stability was 0.52 for both subgroups, and the bootstrap-resampled partition reproduced the original with a mean adjusted Rand index of 0.61 ± 0.48(per-cluster Jaccard distributions in Supplementary Figure S11 and Supplementary Table S14). Meanwhile, the partition was fully recovered when clustering was performed on reduced dimensions (k-means on the first 2, 3, 5, or 10 principal components each gave ARI = 1.000 versus the full 87-feature partition), indicating that the split is not an artifact of high feature dimensionality relative to sample size. Together these analyses indicate that Type 1-A and Type 1-B are best interpreted as two reproducibly identifiable regions along a continuous approach-zone charge axis rather than as discretely separated clusters; The two clusters separated primarily along PC2 in PCA space (Figure 2A). Two outliers at PC1 approximately 15-17 were far removed from the main cluster and may represent sequences with extreme compositions.

**Figure 2.**
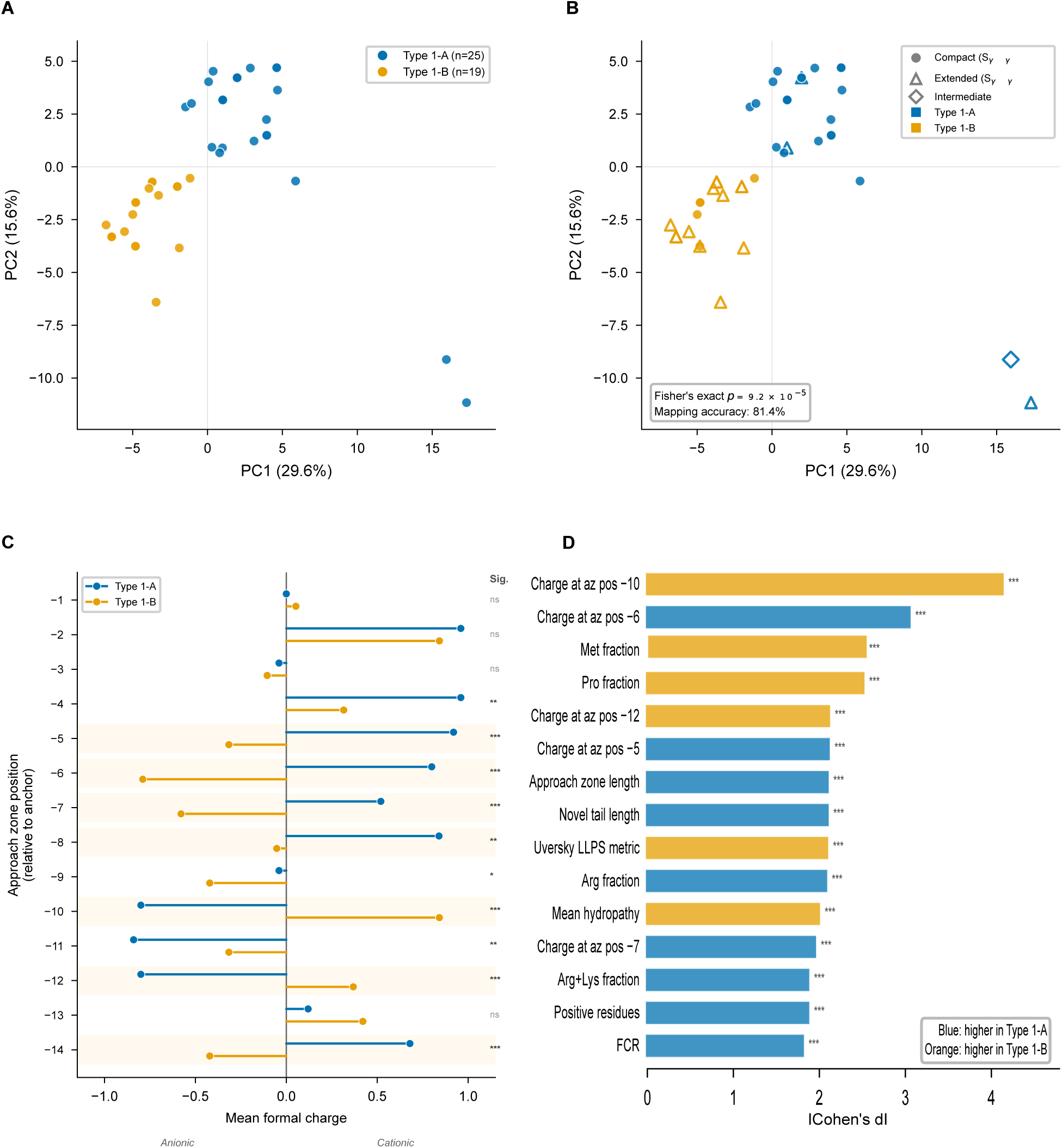
Unsupervised clustering of Type 1-like variants, approach zone charge divergence, and post-hoc feature importance. (A) Principal component analysis (PCA) of 44 Type 1-like variants on 87 standardized sequence features, colored by sequence-derived cluster membership (Type 1-A, blue circles; Type 1-B, orange squares). PC1 accounts for 29.6% and PC2 for 15.6% of total variance; the first two components together explain 45.2%. Type 1-A clusters predominantly in positive PC2 space; Type 1-B in negative PC2 space, with partial overlap near PC2 = 0. Two outlier variants at PC1 ≈ 16–17 are shown at their computed positions. (B) Principal-component projection coloured by predicted Sγ–Sγ geometry. In the rank-1 models, sequence-derived clusters correspond to geometry group (Fisher’s exact p = 9.2 × 10⁻⁵; mapping accuracy 35/43 = 81.4%, one intermediate-geometry variant excluded). This correspondence is specific to the rank-1 model and falls to 36.4% across five models per variant (see Figure 5G). Sγ–Sγ distance was not a clustering input. (C) Approach zone charge profiles for Type 1-A (blue) and Type 1-B (orange). Each horizontal bar shows the mean formal charge at each approach zone position relative to the anchor (position −1 immediately N-terminal to the anchor). Shaded rows indicate positions reaching statistical significance by BH-corrected Mann-Whitney U test. Type 1-A displays a predominantly cationic approach zone; Type 1-B displays anionic character at positions −5, −6, −10, and −14. Significance levels: *p < 0.05; **p < 0.01; ***p < 0.001; ns, not significant. (D) Post-hoc feature importance: top 15 sequence features discriminating Type 1-A from Type 1-B by absolute Cohen’s d effect size, from BH-corrected Mann-Whitney U analysis across 83 tested features. Blue bars indicate features higher in Type 1-A; orange bars indicate features higher in Type 1-B. The dominant discriminators are formal charge at approach zone position −10 (|d| > 4.0, higher in Type 1-B) and position −6 (|d| > 3.5, higher in Type 1-B). All 15 features shown are ***p < 0.001 after BH correction.

### Approach zone charge polarity reversal is the dominant axis of within-Type 1 variation

Post-hoc feature importance analysis identified 50 of 83 tested features as BH-significant discriminators between Type 1-A and Type 1-B (full ranked results in Supplementary Table S8). The absolute Cohen’s d effect sizes are presented in Figure 2D The single strongest discriminating feature was the formal charge at approach zone position -10, followed by approach zone position -6. The approach zone charge profiles (Figure 2C) revealed the mechanistic basis for these effect sizes. Type 1-A displayed a strongly cationic approach zone, with mean formal charge approaching +1.0 at positions -4, -7, -8, and -12. Type 1-B displayed a strikingly different profile, with anionic character (mean charge approximately -0.5 to -1.0) at positions -5, -6, -10, and -14. Statistical comparison at each position showed that divergence was greatest at positions -5 through -8, -10 through -12, and -14 (all p < 0.01 to p < 0.001), while positions -1 through -3 and -13 were not significantly different. This charge polarity reversal at specific approach zone positions constitutes the dominant axis of within-Type 1 sequence variation. Beyond approach zone charges, features enriched in Type 1-B include Methionine fraction, Proline fraction, and approach zone length. Features enriched in Type 1-A included FCR, positive residue count, Arg+Lys fraction, Arg fraction, mean hydropathy, and the charge–hydropathy (CH) disorder-boundary metric. Type 1-A was additionally enriched for RRR triple-arginine motifs (; Supplementary Figure S5A). Three-group pairwise comparisons confirmed that Type 1-A and Type 1-B were each distinguishable from Type 2 (65/88 and 60/84 BH-significant features) and from each other (48/84). Consistently with this approach-zone-driven divergence, wild-type C-domain fragment length did not differ significantly between the two subgroups, confirming that all biophysical divergence between them originates from the approach zone (Figure 1C). Next, we characterized how all three variant classes (Type 1-A, Type 1-B, and Type 2) differ in gross sequence properties, charge architecture, and electrostatic properties.

### Novel tail length distinguishes all three variant groups

Novel tail length differed significantly across the three computationally defined variant groups (Figure 1B). Type 2 variants had the longest novel tails with strikingly low variance, followed by Type 1-A and Type 1-B. All pairwise contrasts were highly significant after BH correction. The approximately 18-residue difference between the shortest (Type 1-B) and longest (Type 2) novel tails is structurally substantial: an additional 18 residues in a disordered tail domain contributes approximately 50-60 Angstrom of additional contour length, expanding the spatial volume sampled by these tails and their capacity to potentially engage distant interaction partners.

### Charge architecture defines three distinct electrostatic regimes

The three variant groups exhibited profoundly different charge architectures in their novel C-terminal tails, with all tested charge features reaching high significance across pairwise comparisons (Figure 3A-E; complete pairwise statistics for all sequence features in Supplementary Table S6). Net charge per residue (NCPR) formed a clear descending hierarchy: Type 1-A, Type 1-B, and Type 2 (, respectively (Figure 3A). Type 1-A tails carry the most pronounced excess positive charge.

**Figure 3.**
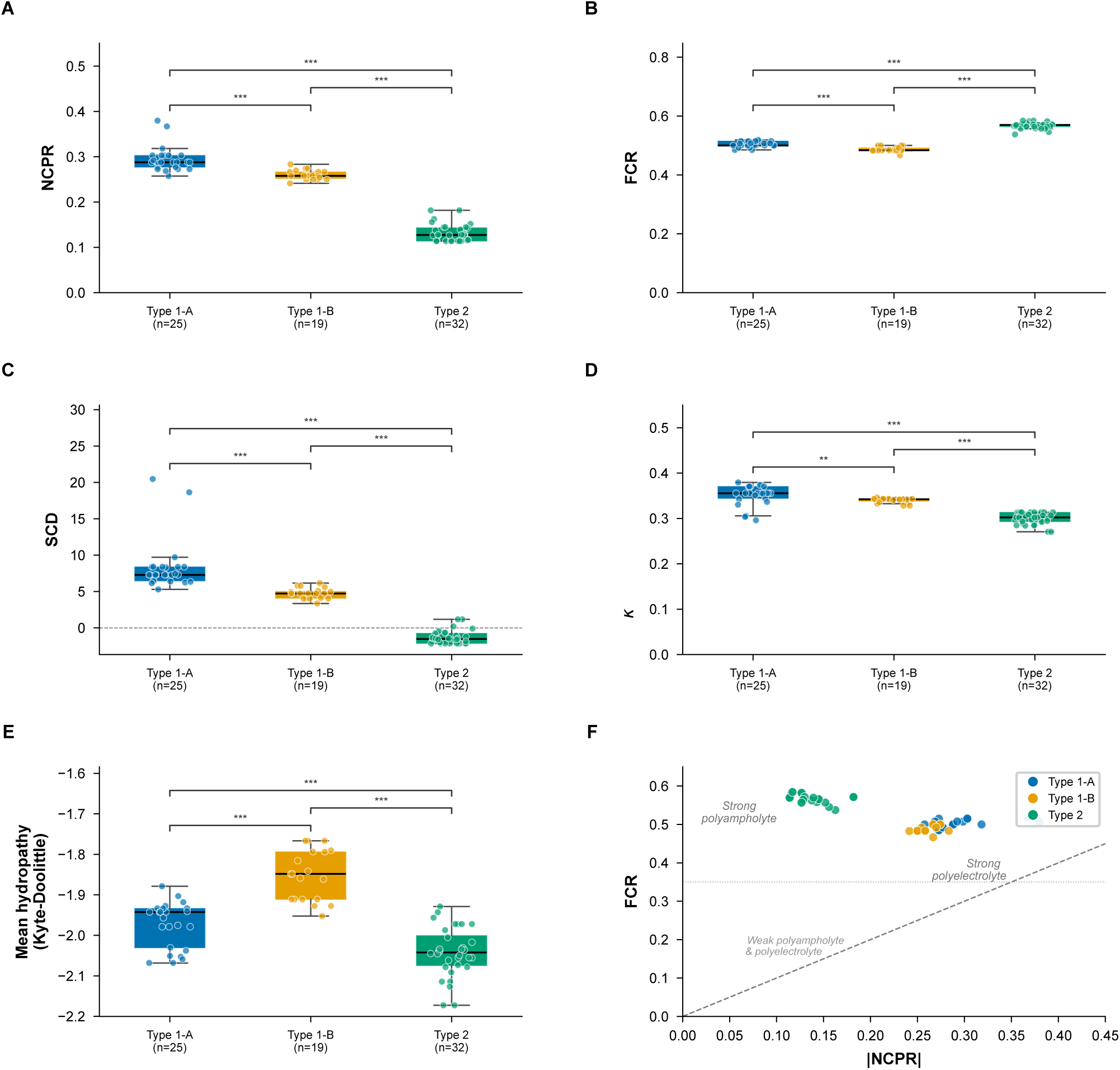
Charge architecture of novel C-terminal tails across the three variant groups. (A) Net charge per residue (NCPR). Type 1-A highest (mean 0.293 ± 0.028), Type 1-B intermediate (0.258 ± 0.019), Type 2 lowest (0.132 ± 0.019); all pairwise ***p < 0.001. (B) Fraction of charged residues (FCR): Type 2 highest (0.567 ± 0.010), Type 1-A intermediate (0.504 ± 0.009), Type 1-B lowest (0.487 ± 0.008); all pairwise p < 0.001. (C) Sequence charge decoration (SCD), quantifying spatial charge patterning. Type 1-A strongly positive (mean 8.32 ± 3.53, charge block segregation), Type 1-B intermediate positive (4.75 ± 0.73), Type 2 weakly negative (mean −1.30 ± 0.92, charge mixing); all pairwise ***p < 0.001. Dashed line at SCD = 0. (D) Charge segregation parameter κ (kappa). Type 1-A highest (0.352 ± 0.022), Type 1-B intermediate (0.340 ± 0.006), Type 2 lowest (0.300 ± 0.013); Type 1-A vs Type 1-B **p < 0.01; all other pairwise ***p < 0.001. (E) Mean Kyte-Doolittle hydropathy of the novel tail. Type 1-B least hydrophilic (mean −1.853 ± 0.062); Type 1-A (−1.972 ± 0.056) and Type 2 (−2.042 ± 0.061) both more hydrophilic; all pairwise ***p < 0.001. Kruskal-Wallis H = 55.83, p < 10⁻¹². (F) Das–Pappu diagram (|NCPR| vs FCR). By the localCIDER phase-plot boundaries all three groups classify as strong polyampholytes (region 3); Type 1-A and Type 1-B lie at the high-positive-charge edge approaching the cationic strong-polyelectrolyte boundary, while Type 2 lies more centrally.

The fraction of charged residues (FCR) revealed an inverted hierarchy relative to NCPR: Type 2 exhibited the highest FCR, followed by Type 1-A and Type 1-B, all pairwise p < 0.001 (Figure 3B). This dissociation between NCPR and FCR is a central finding: Type 2 tails possess the lowest net charge but the highest total charge density, indicating roughly equal proportions of positively and negatively charged residues. This polyampholytic character creates abundant intramolecular salt-bridge opportunities and intermolecular electrostatic interaction sites that differ fundamentally from the charge-excess-driven interactions expected for Type 1 tails.

SCD, which quantifies the spatial patterning of charged residues along the primary sequence, provided the most dramatic group separation (Figure 3C). Type 1-A displayed strongly positive SCD values, Type 1-B showed intermediate positive values, and Type 2 fell into weakly negative territory. Positive SCD indicates that like-charged residues are spatially clustered into blocks, whereas negative SCD reflects an alternating or interspersed charge distribution. The charge segregation parameter kappa corroborated this pattern, with Type 1-A highest, Type 1-B intermediate, and Type 2 lowest (Figure 3D). The Type 1-A versus Type 1-B comparison for kappa reached only p < 0.01, compared to p < 0.001 for all other pairwise tests, suggesting that the magnitude of charge patterning (SCD) differs more dramatically than the normalized topology of charge segregation (kappa) between the Type 1 subtypes. Because κ is less reliable for sequences with very low negative-charge content, the κ values for Type 1-A should be interpreted with caution: 3 of 25 Type 1-A variants have f⁻ < 0.10, where the κ patterning parameter becomes poorly defined.

### Sequence-based ensemble dimensions support the charge-regime interpretation

Consistent with the charge-regime analysis, ALBATROSS predicted all three groups to be disordered, with apparent scaling exponents (ν) of 0.49 (Type 2), 0.54 (Type 1-A), and 0.54 (Type 1-B), and predicted radii of gyration of 25.9Å (Type 2), 26.8 Å (Type 1-A), and 24.7 Å (Type 1-B). The inter-group diffcounterintuitive;a more expanded predicted monomer tail coincides with the more compact predicted dimer class,andetrics broadly corroborate the disordered, expanded nature of all three tails rather than sharply distinguishing them. Within the Type-1 group, predicted monomer Rg was inversely correlated with the predicted inter-chain Sγ–Sγ distance, whereas no such correlation was present in Type 2. The inverse sign is counterintuitive;a more expanded predicted monomer tail coincides with the more compact predicted dimer class,and is consistent with the compact Type 1-A dimers being held together by post-anchor inter-chain contacts We present this as an observation requiring experimental testing rather than an established relationship (Supplementary Table S12). Mean Kyte-Doolittle hydropathy differed significantly across the three variant groups (Figure 3E). Type 2 novel tails were the most hydrophilic, Type 1-A was similarly hydrophilic, and Type 1-B was significantly less hydrophilic.

The Das-Pappu charge classification diagram (Figure 3F) provided an integrative synthesis. By the formal Das-Pappu boundaries (localCIDER phase-plot.), all three groups classify as strong polyampholytes (region 3), reflecting their high overall fraction of charged residues. They are distinguished by their position within the region: Type 2 lies centrally in the strong-polyampholyte regime, where charge-balanced sequences are predicted to adopt comparatively compact disordered conformations, whereas Type 1-A and Type 1-B lie at the high-positive-charge edge of the region, approaching the cationic strong-polyelectrolyte boundary. This net-cationic character is predicted to favour more expanded conformations for the Type 1 tails. The three groups showed minimal overlap in this charge space, confirming that a simple two-parameter classification captures the dominant electrostatic variation.

To place the groups in the Das–Pappu diagram of states using its native coordinates, we computed the fraction of positive (f⁺) and negative (f⁻) residues per group and assigned phase-plot regions. All three groups fall predominantly in the strong-polyampholyte region (region 3) The Type 1 subgroups lie at the high-f⁺ edge of the strong-polyampholyte region, approaching the cationic polyelectrolyte boundary, consistent with their high net positive charge, whereas Type 2 lies more centrally owing to its higher negative-charge fraction (Supplementary Table S13).

### Polyanion runs, charge contrast, and polycation runs define type-specific electrostatic strategies

The longest polyanion run showed a dramatic Type 2-specific signature (Figure 4A). The longest polycation both centered at approximately 3 residues with no significant difference between them, while Type 2 exhibited an invariant longest polyanion run of exactly 5 residues across all 32 sequences. The wild-type-novel charge contrast showed a clear three-group hierarchy (Figure 4B), quantifying the electrostatic asymmetry between the anionic conserved C-domain and the cationic novel tail, and directly predicts the strength of heterotypic inter-chain attraction in the homodimer (see Discussion).

**Figure 4.**
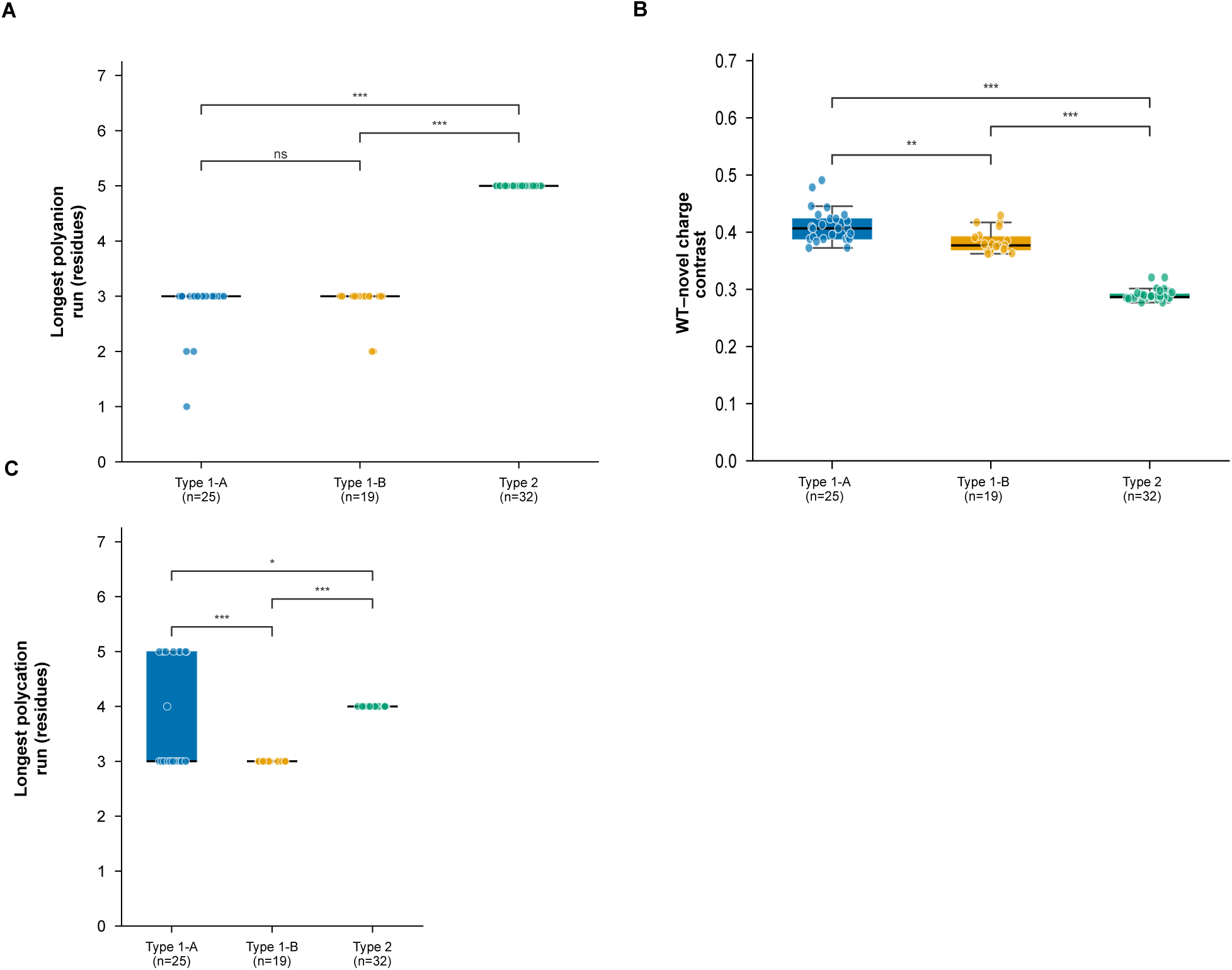
Type-specific electrostatic strategies: polyanion architecture, charge contrast, and polycation runs. (A) Longest consecutive polyanion run (residues). Type 2 exhibits an invariant run of exactly 5 residues in all 32 variants (mean 5.000 ± 0.000); Type 1-A (mean 2.84 ± 0.47) and Type 1-B (2.89 ± 0.32) both center at 3 residues with no significant difference between them; Kruskal-Wallis H = 62.85, p = 8.40 × 10⁻¹⁶, ε² = 0.834. (B) Wild-type C-domain to novel tail charge contrast. Type 1-A highest (mean 0.411 ± 0.030), Type 1-B intermediate (0.384 ± 0.021), Type 2 lowest (0.290 ± 0.010); all pairwise **p < 0.01 to ***p < 0.001. This metric quantifies the electrostatic asymmetry between the anionic conserved C-domain and the cationic novel tail. (C) Longest consecutive polycation run (residues). Type 1-B invariant at exactly 3 in all 19 variants (mean 3.000 ± 0.000); Type 2 invariant at exactly 4 in all 32 variants (mean 4.000 ± 0.000); Type 1-A variable and highest (mean 3.680 ± 0.945, median 3); Type 1-A vs Type 1-B ***p < 0.001; Type 1-A vs Type 2 *p < 0.05; Type 1-B vs Type 2 ***p < 0.001. Box plots: median and IQR; whiskers 1.5× IQR; individual points shown. *p < 0.05; **p < 0.01; ***p < 0.001; ns, not significant (BH-corrected Mann-Whitney U).

We quantified the acidic wild-type C-domain charge retained upstream of each frameshift, since the heterotypic-attraction model requires that an anionic partner surface remains. All three groups retain a substantial portion of the native acidic C-domain.; the del52 frameshift removes roughly 44% of the native acidic C-domain rather than all of it. The anionic partner surface required by the heterotypic-attraction model is therefore present in all groups, and the inter-group differences in predicted dimer geometry are driven by the cationic novel tail rather than by loss of the acidic domain.

The longest polycation run differed significantly across the three variant groups: Type 1-B invariant at exactly 3 residues, Type 2 invariant at exactly 4 residues, and Type 1-A variable and highest (Figure 4C).

### Structural predictions are concordant with sequence subgroups at the Type level

ColabFold v1.5 homodimer structural predictions were analyzed as an independent post-hoc validation of the sequence-derived subgroup classification, as structural features were not used as clustering inputs (complete pairwise structural statistics in Supplementary Table S7). Monomer pLDDT scores were high across all groups for the wild-type CALR region (Supplementary Figure S7B). Novel tail pLDDT was moderate in all groups, spanning a wide range (43.4-78.8 overall; Supplementary Figure S7C), with Type 1-B variants frequently falling below pLDDT=50. This moderate confidence may reflect the anchor region providing a structured scaffold that elevates per-residue confidence throughout the novel tail, or may represent an AlphaFold2 artifact in the absence of coevolutionary signal for these frameshift-derived sequences (see Discussion).

### Bimodal cysteine geometry maps onto sequence-derived clusters

The most structurally discriminating feature across groups was the Sγ–Sγ inter-chain distance at the anchor+32 cysteine (Figure 5A). This distance was bimodally distributed across the 44 Type 1-like variants, resolving into a compact conformationand an extended conformation, with one variant at an intermediate distance. The magnitude of the extended distances (140-180 Angstrom) is consistent with predicted models in which in extended dimers, the novel tails project outward in essentially opposite directions without making significant inter-chain contacts. Compact distances of 8-15 Angstrom place the cysteines in close spatial proximity, consistent with intimate tail-tail interaction at the dimer interface.

**Figure 5.**
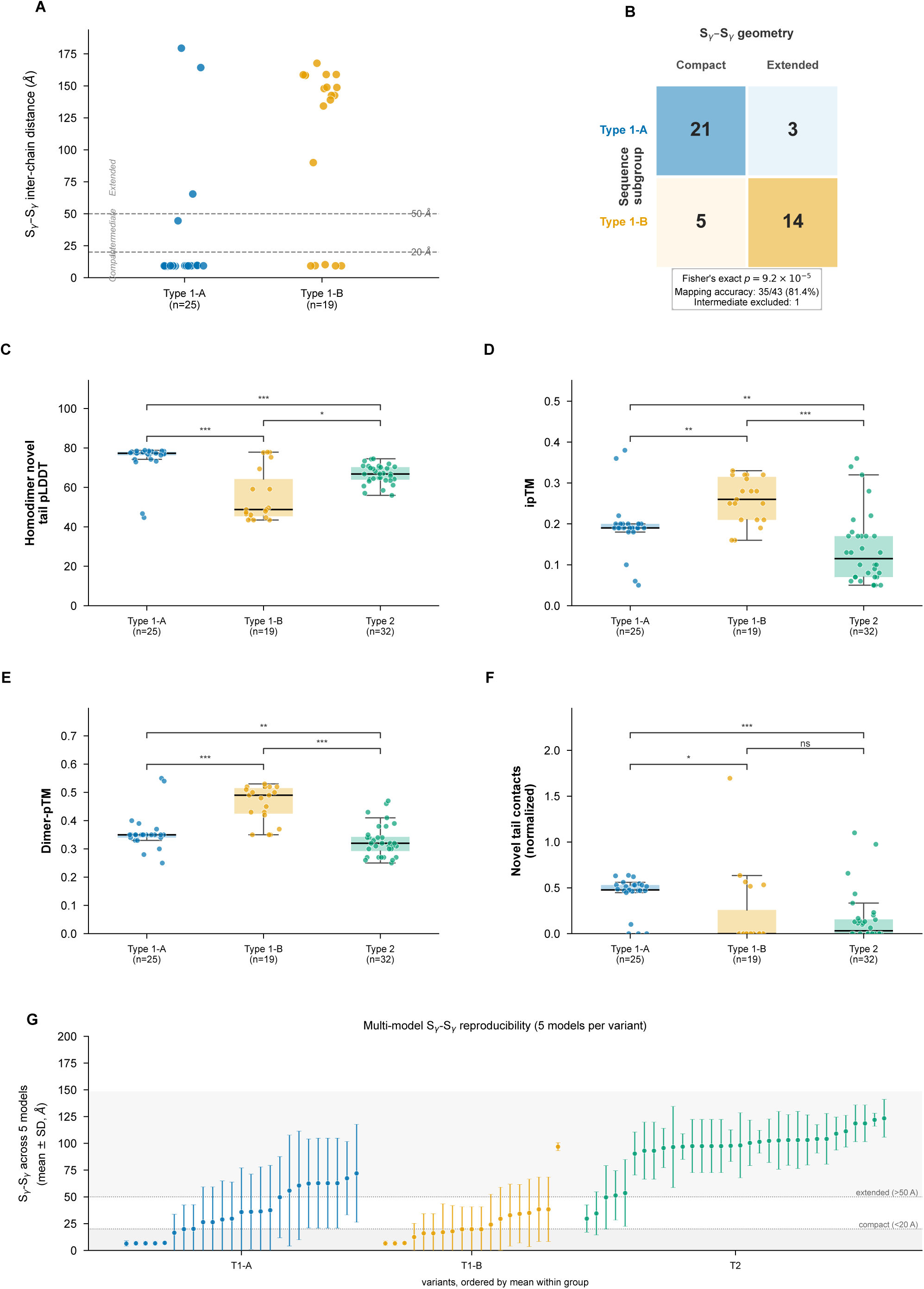
ColabFold homodimer structural predictions and their concordance with sequence-derived subgroups. (A) Sγ–Sγ inter-chain distance (Å) at the anchor+32 cysteine for all 44 Type 1-like variants, from unrelaxed rank-1 ColabFold v1.5 homodimer models. Dashed lines mark the 20 Å and 50 Å thresholds defining compact (< 20 Å, n = 26) and extended (> 50 Å, n = 17) geometry groups; one variant at ∼44.5 Å is classified as intermediate. Type 1-A is predominantly compact (21/25 variants); Type 1-B predominantly extended (14/19 variants). (B) Contingency heatmap of sequence subgroup versus Sγ–Sγ geometry group. Cell values indicate variant counts. Rank-1 models: Fisher’s exact p = 9.2 × 10⁻⁵; mapping accuracy 35/43 = 81.4%; one intermediate variant excluded. Across five models per variant this concordance falls to 36.4% (see panel G and Results). (C) Homodimer novel tail pLDDT across all three variant groups. Type 1-A highest (mean 74.46 ± 8.82), Type 1-B lowest (55.30 ± 13.32), Type 2 intermediate (66.67 ± 4.78); Kruskal-Wallis H = 32.29, p = 9.72 × 10⁻⁸, ε² = 0.415. (D) Interface predicted TM-score (ipTM). Type 1-B highest (mean 0.258 ± 0.056), Type 1-A intermediate (0.193 ± 0.055), Type 2 lowest (0.139 ± 0.088). The counterintuitive ordering reflects the dominance of well-folded conserved domain residues in this composite score when disordered tails are extended away from the interface. (E) Dimer predicted TM-score (dimer-pTM). Type 1-B highest (0.460 ± 0.065), Type 1-A intermediate (0.347 ± 0.040), Type 2 lowest (0.326 ± 0.059). (F) Normalized inter-chain novel tail contacts (Cα–Cα < 8 Å, normalized by tail length). Type 1-A shows the highest contact density; Type 1-B and Type 2 substantially lower. Box plots: median and IQR; whiskers 1.5× IQR; individual points shown. *p < 0.05; **p < 0.01; ***p < 0.001; ns, not significant (BH-corrected Mann-Whitney U). Panels A, B, and F report rank-1 model values. (G) Per-variant inter-chain Sγ–Sγ distance (mean ± SD across five ColabFold models), variants ordered by mean within each group (Type 1-A blue, Type 1-B orange, Type 2 teal). Shaded bands mark compact (<20 Å) and extended (>50 Å) regimes. Type 2 is reproducibly extended with small inter-model dispersion; Type 1-A and Type 1-B span the full range with large dispersion and substantial overlap. The compact/extended call is stable across all five models for 35/76 variants; the Type 1-A/Type 1-B subgroups are not separable by mean Sγ–Sγ (Mann–Whitney p = 0.079); rank-1 cluster–geometry concordance (81.4%) falls to 36.4% across five models.

In the rank-1 models, sequence-derived cluster membership corresponded to Sγ–Sγ geometry group with 35/43 = 81.4% accuracy (Figures 2B, 5B). This correspondence is, however, specific to the rank-1 model. Across five independent ColabFold models per variant, the compact/extended assignment was stable for only 35 of 76 variants overall, and within the Type-1 group the 1-A and 1-B subgroups were not separable by mean Sγ–Sγ distance. Concordance between the sequence subgroups and the predicted geometry fell from 35/44 = 79.5% in the rank-1 models to 16/44 = 36.4% at the five-model level (i.e. to chance). The compact/extended distinction is thus robust between Type 1 and Type 2 but does not independently corroborate the Type 1-A/Type 1-B subdivision (see Multi-model analysis below and Limitations).

### Multi-model analysis defines the reproducibility of the predicted geometry

To assess whether the compact/extended assignment depends on the choice of a single (rank-1) model, we generated five ColabFold homodimer models for each of the 76 variants and re-extracted the inter-chain Sγ–Sγ distance from all models. The five models agreed on the relative tail geometry of each variant. The discrete compact/extended call was stable across all five models for 35 of 76 variants. Predicted interface confidence was uniformly low across all variants.Although the predicted geometry was reproducibly governed by approach-zone length, the Type 1-A versus Type 1-B subdivision was not separable by mean Sγ–Sγ across models, and the rank-1 cluster–geometry concordance (81.4%) was not reproduced at the five-model level (36.4%). The predicted geometry therefore provides continuous, reproducible structural correlates of the sequence variation at the Type level, but does not define discrete structural classes within Type 1 (Supplementary Table S11; Figure 5G).

### Confidence metrics reveal a paradox between local structure and global interface prediction

Homodimer novel tail pLDDT differed dramatically across the three variant classes (Figure 5C). Type 1-A showed the highest confidence, Type 1-B showed substantially lower values, and Type 2 was intermediate.

A counterintuitive pattern emerged from the interface confidence metrics (Figure 5D-E). ipTM was highest for Type 1-B, intermediate for Type 1-A, and lowest for Type 2. Dimer-pTM mirrored this pattern: Type 1-B highest Type 1-A intermediate, Type 2 lowest.

This paradox is resolved by the compositional nature of these metrics: global ipTM is essentially uncorrelated with the tail Sγ–Sγ geometry (Spearman rho = 0.06, n.s.), whereas tail-restricted confidence tracks it strongly. The high global ipTM of the most-extended group therefore reflects confident conserved-domain packing rather than a confident tail interface (Supplementary Figure S12) (see Discussion).

### Contact topology reveals qualitatively distinct homodimer interface architectures

To characterize the spatial organization of the predicted homodimer interface, we extracted all inter-chain Ca-Ca contacts (< 8.0 Angstrom) within the novel C-terminal tailfor all 76 variants. Contacts were classified by the region of origin of each contacting residue.

Contact region analysis revealed fundamentally distinct interface architectures across the three variant groups (Figure 5F; Supplementary Table S10). In Type 1-A homodimers, 88% exhibited at least one inter-chain novel tail contact, yielding 723 total contacts. Of these, 715 (98.9%) occurred between post-anchor regions of opposing chains, with only 1.1% contacts involving approach zone residues. The invariant cysteine residues at anchor-relative positions +32 and +36 constituted the dominant contact hotspots. This finding is consistent with the compact Sγ–Sγ geometry that predominates in Type 1-A and demonstrates that the predicted homodimer interface in this subgroup is organized around inter-chain cysteine proximity in the post-anchor region.

In striking contrast, Type 2 homodimers showed a fundamentally different contact topology. Only 50% had any inter-chain contacts, yielding 398 total contacts. Of these,55.5%occurred between approach zones of opposing chains, and an additional27.1% involved anchor-approach zone cross-contacts, such that 82.6% of all Type 2 contacts involved the approach zone. Only 11.6% were post-anchor-post-anchor. The most frequently contacted position was approach zone position -1 (53 contacts, of which 51 involved cysteine residues), followed by position -4. In the predicted Type-2 homodimers that formed a defined interface, the two identical chains paired in an antiparallel register, with the anchor of one chain contacting the post-anchor of the other in a symmetric, self-complementary arrangement. The most extended Type-2 predictions formed minimal interface. This register is consistent with the charge-alternating, self-complementary pairing proposed above, though it is derived from low-confidence predicted models and requires experimental testing.

Type 1-B homodimers exhibited a sparse, mixed contact pattern. Only 26% exhibited inter-chain contacts, yielding 239 total contacts. Of these,66.9%were post-anchor-post-anchor and24.7% were approach-approach, reflecting the intermediate structural character of this subgroup. The high variance and median of 0 indicate that a few Type 1-B variants make substantial contacts while the majority make none.

Across all three groups, contacts were predominantly cross-chain (different positions on the two chains contacting each other) rather than diagonal (equivalent positions): Type 1-A 91.0% cross, Type 1-B 95.0% cross, Type 2 91.5% cross.

### Inter-chain distance profiles confirm region-specific proximity patterns

Mean inter-chain Ca-Ca distances at equivalent positions quantified the spatial separation (Supplementary Figure S10) between the two chains along the full length of the novel tail. In the approach zone, Type 2 showed the shortest inter-chain distances across all positions, Type 1-A was intermediate, and Type 1-B was similar to Type 1-A. In the post-anchor region, the pattern inverted dramatically: Type 1-A showed tight inter-chain proximity, with the shortest distances at the cysteine positions +32and +36, while Type 1-B and Type 2 showed widely separated post-anchor chains.

This reciprocal distance pattern confirms the contact topology: Type 1-A brings its post-anchor regions into proximity while approach zones remain apart, whereas Type 2 brings its approach zones into proximity while post-anchor regions remain distant. Type 1-B achieves proximity in neither region efficiently.

## Discussion

This study demonstrates that the physicochemical, predicted structural, and interface topological diversity of CALR novel C-terminal tails extends substantially beyond the canonical classification. Through systematic analysis of sequence features across 76 CALR mutants we identify three computationally distinct variant classes, Type 2, Type 1-A, and Type 1-B, each exhibiting characteristic electrostatic signatures, charge patterning regimes, predicted structural profiles, and qualitatively distinct homodimer interface architectures. The central finding is that the three variant groups do not merely differ in the magnitude of shared properties but may be employing fundamentally different molecular mechanisms for inter-chain engagement: Type 1-A through cysteine-proximal post-anchor contacts, Type 2 through approach zone charge-complementary contacts, and Type 1-B through a weakly engaged mixed interface.

### Charge architecture and the Das-Pappu framework

The charge architecture differences across the three variant groups recapitulate and substantially extend known biology. The high cationic content of Type 1-like novel tails relative to Type 2is consistent with the established requirement for positive charge in the novel tail for TpoR engagement (5,10,24). The SCD analysis adds a dimension not previously quantified across a large variant panel: the distinction between charge segregation (positive SCD in Type 1) and charge mixing (negative SCD in Type 2) represents a fundamental difference in charge topology that has direct structural consequences demonstrated by our contact topology analysis (see below). The FCR analysis places the three variant groups in distinct regimes of the Das-Pappu IDP classification framework: Type 2 in the strong polyampholyte zone (compact globular conformations), Type 1-A extending into the strong polyelectrolyte zone (extended swollen conformations), and Type 1-B straddling the boundary. This classification predicts distinct conformational ensembles for the three tail types that could be tested by solution-state biophysical methods such as small-angle X-ray scattering or single-molecule FRET.

The perfect co-occurrence of the KKRK motif with Type 2 classification (32/32), combined with the invariant presence of exactly three consecutive acidic clusters, exactly two KEE motifs, and polyanion runs of exactly 5 residues in all 32 Type 2 variants, defines an extraordinarily constrained sequence architecture suggesting strong selective constraint on the Type 2 tail in MPN, consistent with its association with a distinct clinical phenotype (8). Type 2 variants also retained substantially more wild-type C-domain sequence, potentially preserving native calcium-binding function of the CALR acidic C-domain (6,25).

### Approach zone charge architecture as a novel discriminating axis

The identification of two sequence-driven subgroups within Type 1-like variants is a central novel finding. Both clustering algorithms recovered the same partition, and sensitivity analyses demonstrated robustness to imputation and feature redundancy (ARI = 1.000), although bootstrap resampling showed the membership boundary itself is soft. To address the concern that defining the subgroups partly on approach-zone charge features and then identifying those same features as discriminators is circular, we note that the approach zone is the only sequence-variable region across these variants. The subgroups are therefore, by construction, a partition of approach-zone sequence space, and excluding the approach zone entirely is uninformative rather than a meaningful control. We do not claim independent feature-based corroboration of the partition; its reproducibility is established by the algorithm concordance and dimension-robustness above, and its honest status as a soft-bounded partition by the bootstrap analysis. The dominant discriminating axis is the charge polarity pattern of the approach zone, with charge polarity reversal at specific positions which have direct structural implications that are revealed by the inter-chain distance data: Type 1-A’s strongly cationic approach zone positions produce potential electrostatic repulsion between the identical cationic surfaces of the two chains, while Type 2’s charge-balanced approach shows charge-complementary for self-pairing (see below).

### Two distinct homodimer interface architectures

The contact topology analysis is the most mechanistically suggestive finding of this study. In the rank-1 predicted models, the contact interfaces are largely segregated; Type 1-A making 98.9% of contacts in the post-anchor region versus Type 2 making 82.6% of contacts involving the approach zone, suggesting that these variant groups may employ qualitatively different dimerization mechanisms rather than differing merely in the efficiency of a shared mechanism (per-variant inter-chain contact pairs in Supplementary Table S9). Because these patterns derive from single (rank-1) predicted structures with low predicted interface confidence (see Limitations), they are best regarded as hypotheses for experimental testing. We note that this contact-topology comparison contrasts Type 1-A with Type 2 and therefore was not subjected to the five-model stability check applied to the within-Type-1 Sγ–Sγ distinction; it is reported from the rank-1 models as a mechanistic hypothesis. In Type 1-A homodimers, the interface is organized around the invariant post-anchor cysteines at positions +32 and +36.Because the post-anchor region is invariant across all variants, the Type 1-A interface utilizes the conserved structural elements shared by all CALR frameshift variants. The question then becomes: why does Type 1-A, but not Type 1-B or Type 2, achieve post-anchor cysteine proximity? The answer lies in the approach zone charge environment. Type 1-A’s high NCPR high positive SCD, and RRR motifs create strong heterotypic electrostatic attraction between the cationic novel tail of one chain and the anionic conserved C-domain of the opposing chain. The WT-novel charge contrast, which is highest in Type 1-A directly quantifies this electrostatic asymmetry. This heterotypic attraction may bring the post-anchor regions of the two chains into proximity, positioning the cysteines within distances compatible with disulfide bond formation. Once formed, the disulfide bonds lock the compact dimer state.

This interpretation is independently supported by molecular dynamics simulations of CALR C-domain dimers by Radjasandirane and de Brevern (2023) (23). Using GROMACS MD with the CHARMM-36 forcefield, they demonstrated that class A (canonical Type 1) dimers with intact disulfide bonds remained stable over 500 nano second of simulation, but that positive electrostatic patches from the middle of both chains induced repulsive forces, producing inter-chain distances of 25-85 Angstrom in regions distal to the disulfide bonds. When the disulfide bridge was reduced, the two chains repelled each other immediately, with distances exceeding 100 Angstrom within nanoseconds. This directly supports our observation that Type 1-like tails carry high positive NCPR, creating identical cationic surfaces that generate inter-chain repulsion that must be overcome by the heterotypic attraction and then locked by disulfide bonds.

In Type 2 homodimers, the interface is instead organized around the approach zone. This approach-zone-mediated interface architecture is consistent with Type 2’s charge properties: the negative SCD indicates alternating positive and negative charges along the sequence, creating a charge pattern that is self-complementary where when two identical Type 2 approach zones face each other, positive residues on one chain can align with negative residues on the opposing chain in a charge-complementary register, forming intermolecular salt-bridge networks. This mechanism does not require net charge asymmetry; it requires only charge alternation, which the SCD metric specifically quantifies.

Radjasandirane and de Brevern (2023) provide indirect support for this distinct Type 2 interface. They showed that class B (canonical Type 2) monomers adopt a three-helix architecture with distinct dynamics from the single-helix class A (canonical Type 1), and that class B dimers showed qualitatively different distance dynamics from class A dimers. While their study examined only one Type 2 representative (Ins5), our population-level analysis across 32 Type 2 variants reveals that the approach-zone-mediated interface is a consistent feature of the Type 2 class, not an idiosyncrasy of the canonical variant.

Type 1-B homodimers display a sparse, mixed contact pattern which is consistent with Type 1-B’s intermediate charge properties: the approach zone contains anionic positions that partially support charge-complementary pairing but without the consistent alternating pattern of Type 2, while the overall charge driving force is insufficient for robust cysteine proximity in the post-anchor region. The high variance in contacts per variant indicates that a few Type 1-B variants with borderline charge properties can engage one or other mechanism, while the majority engage neither efficiently.

### The heterotypic attraction model

The contact topology data resolves an apparent paradox in the charge-geometry relationship. Naively, one might predict that variants with the lowest inter-chain electrostatic repulsion (lowest NCPR: Type 2) should show the most compact homodimer geometry. The opposite is observed: Type 2 shows the longest post-anchor inter-chain distances despite having the least positive charge. This paradox is resolved by recognizing that the determinant of post-anchor cysteine proximity is not the magnitude of inter-chain repulsion but rather the magnitude of heterotypic inter-chain attraction between the cationic novel tail of one chain and the anionic conserved C-domain of the opposing chain.

Type 1-A’s high NCPR, high positive SCD, RRR motifs, and high WT-novel charge contrast may plausibly create strong heterotypic attraction that overcomes the homo-charge repulsion between the two tails, driving the post-anchor cysteines together. Type 1-B’s weaker charge properties produce insufficient heterotypic attraction, and the cysteines never achieve proximity. Type 2’s charge-balanced, polyampholytic tails satisfy their charges intramolecularly through short-range salt bridges, leaving no free cationic charge to engage the anionic partner domain, and the tails remain electrostatically self-contained in the post-anchor region.

However, Type 2 may have an alternative: the approach zone self-complementarity mechanism. Rather than driving cysteine proximity through heterotypic attraction, Type 2 forms an interface through the variant-specific approach zone, utilizing the charge-alternating architecture that is a hallmark of its sequence. This may represent an alternative dimerization mechanism that maintains inter-chain engagement capacity despite the charge-balance properties that preclude post-anchor cysteine proximity. The cysteine at approach zone position -1 in Type 2 variants (contributing 51 of 53 contacts at that position) may serve as an alternative cross-linking point for disulfide-mediated stabilization of the approach-zone interface, though this hypothesis requires experimental confirmation.

### Convergence with molecular dynamics simulations

Our findings converge with and extend the molecular dynamics results of Radjasandirane and de Brevern (2023) (25) in several important aspects. First, their demonstration that positive electrostatic patches produce inter-chain repulsion in Type 1 dimers provides dynamic validation of the charge-based mechanisms we infer from static ColabFold predictions and sequence analysis. Second, their finding that disulfide bonds are essential to maintain the dimeric form of classes A, B, and C is consistent with our observation that the CREAC motif cysteines at positions +32 and +36 constitute the dominant contact hotspots in Type 1-A homodimers (32.1% of all contacts). Third, their note that CALRwt and class E (which lack novel cysteines) cannot dimerize validates the fundamental requirement for cysteine-mediated interactions that all 76 variants in our dataset share through the invariant cysteine residues. Importantly, Radjasandirane and de Brevern also noted that AlphaFold2 and ColabFold produced unsatisfying results for the novel tail region due to empty multiple sequence alignments, because the frameshift-derived sequences do not exist in natural proteomes. This MSA depth limitation applies equally to our ColabFold predictions and underscores that all novel tail structural metrics in both studies are computational hypotheses. The convergence of their MD approach (dynamic but limited to single representatives) with our ColabFold approach (static but covering 76 variants) strengthens confidence in shared conclusions while highlighting the distinct limitations of each method.

Their study examined 6 representative variants (one per class) through deep MD simulation over hundreds of nanoseconds, while our study examines 76 variants through comprehensive sequence analysis and AlphaFold structural prediction. The approaches are complementary: their MD provides dynamic resolution and force-field-based validation that static prediction cannot offer, while our population-level analysis reveals statistical patterns across the variant landscape that single-variant MD cannot capture; most notably, the distinct contact topologies and the approach zone charge divergence between Type 1-A and Type 1-B that would not be apparent from studying only the canonical Type 1 representative.

### Counterintuitive confidence metrics

The counterintuitive confidence metric pattern, highest ipTM and dimer-pTM in Type 1-B despite its extended geometry and low tail pLDDT, provides important methodological insight for interpreting AlphaFold multimer predictions of proteins with intrinsically disordered regions. ipTM and dimer-pTM are composite scores that weight all residue pairs, including well-folded conserved domains. When disordered tails are extended away from the interface (Type 1-B), the conserved domain-domain interface dominates the score. When disordered tails participate in the interface (Type 1-A), they introduce prediction uncertainty that depresses global scores despite high local confidence. This cautions against using ipTM or dimer-pTM as sole arbiters of dimer confidence when IDR-mediated contacts are involved and suggests that region-specific metrics provide more informative readouts.

### An integrated three-strategy model

Synthesizing across all analyses, the data support a model in which the three variant classes employ distinct molecular strategies for homodimer formation. Type 1-A tails function as cationic, charge-segregated effectors that drive heterotypic attraction between the novel tail and the anionic conserved domain of the opposing chain, bringing the post-anchor cysteines into disulfide-bonding proximity and producing a cysteine-nucleated, post-anchor interface. Type 1-B tails are shorter, less charged, enriched in structure-breaking residues (Pro, Met), and generate insufficient heterotypic attraction for efficient cysteine proximity, producing a sparse, weakly engaged interface. Type 2 tails are long, charge-balanced polyampholytes that satisfy their charges intramolecularly in the post-anchor region but achieve inter-chain engagement through charge-complementary self-pairing in the approach zone, producing an approach-zone-mediated interface that does not depend on post-anchor cysteine proximity.

The observation that Type 1-B occupies an intermediate position on virtually every metric raises the question of whether it represents a truly distinct functional class or a transitional phenotype. However, the clustering was reproduced by two algorithms with perfect agreement, and the charge polarity reversal at specific approach zone positions is a qualitative sign change rather than a quantitative magnitude shift. The bimodal Sγ–Sγ distribution seen in the rank-1 models is consistent with this subdivision but is not reproduced across the five-model ensemble (see Results and Limitations); we therefore treat the Type 1-A/Type 1-B distinction as sequence-defined rather than structurally discrete. We propose the three-group taxonomy as an exploratory framework for generating testable hypotheses, while acknowledging that individual variants near the Type 1-A/Type 1-B boundary may exhibit intermediate properties.

### Limitations

Several limitations must be acknowledged. First, all findings are computational. Sequence features do not account for post-translational modifications, cellular context, or interaction partners. ColabFold predictions are not equivalent to experimental structures, and a specific limitation concerns MSA depth: because the novel tail sequences arise from somatic frameshifts and do not exist in natural proteomes, MMseqs2 returns effectively zero homologs for this region, so AF2 predictions rely on single-sequence representations rather than coevolutionary signal. This limitation is shared with the MD study of Radjasandirane and de Brevern (2023), who also noted unsatisfying AlphaFold results for these sequences. All novel tail structural metrics are computational hypotheses requiring experimental validation. The low overall ipTM confirms limited interface confidence, consistent with the IDR nature of the novel tail. A further limitation concerns model dependence. Although primary metrics were extracted from the rank-1 model, we additionally generated five models per variant for all 76 homodimers. The five models agreed on a per-variant basis at the Type level but agreed substantially less within Type 1, and the discrete compact/extended call was stable across all five models for only 35 of 76 variants. Predicted interface confidence was uniformly low. Consequently the rank-1-based Type 1-A/Type 1-B structural correspondence was not reproduced across models, and the Type 1-A/Type 1-B distinction should be regarded as sequence-defined, with predicted geometry providing continuous, model-dependent rather than discrete structural correlates. Second, the dataset of 76 variants remains modest, and Type 1-B has the least statistical power. The weaker significance observed for certain comparisons may partly reflect this power limitation. Third, the PCA explained 45.2% of variance in two components, with nine components required for 80% cumulative variance, indicating substantial remaining dimensionality. Fourth, the contact topology analysis relies on a single distance cutoff (8.0 Angstrom) applied to static ColabFold predictions; the dynamic nature of these interfaces, as demonstrated by Radjasandirane and de Brevern’s MD simulations, means that transient contacts and conformational fluctuations are not captured. Fifth, the subgroups are computationally defined; no clinical outcome data were used, and no inference about differential disease predisposition, thrombotic risk, fibrotic transformation, or treatment response can be drawn. Prospective clinical correlation studies are essential before clinical relevance can be attributed. Lastly, consistent with the weak internal cluster-validity indices, bootstrap resampling showed that the precise membership boundary between Type 1-A and Type 1-B is not robust; we therefore treat the subdivision as an exploratory partition of a continuum rather than as discrete classes.

## SUPPLEMENTARY FIGURES

**Supplementary Figure S1.**
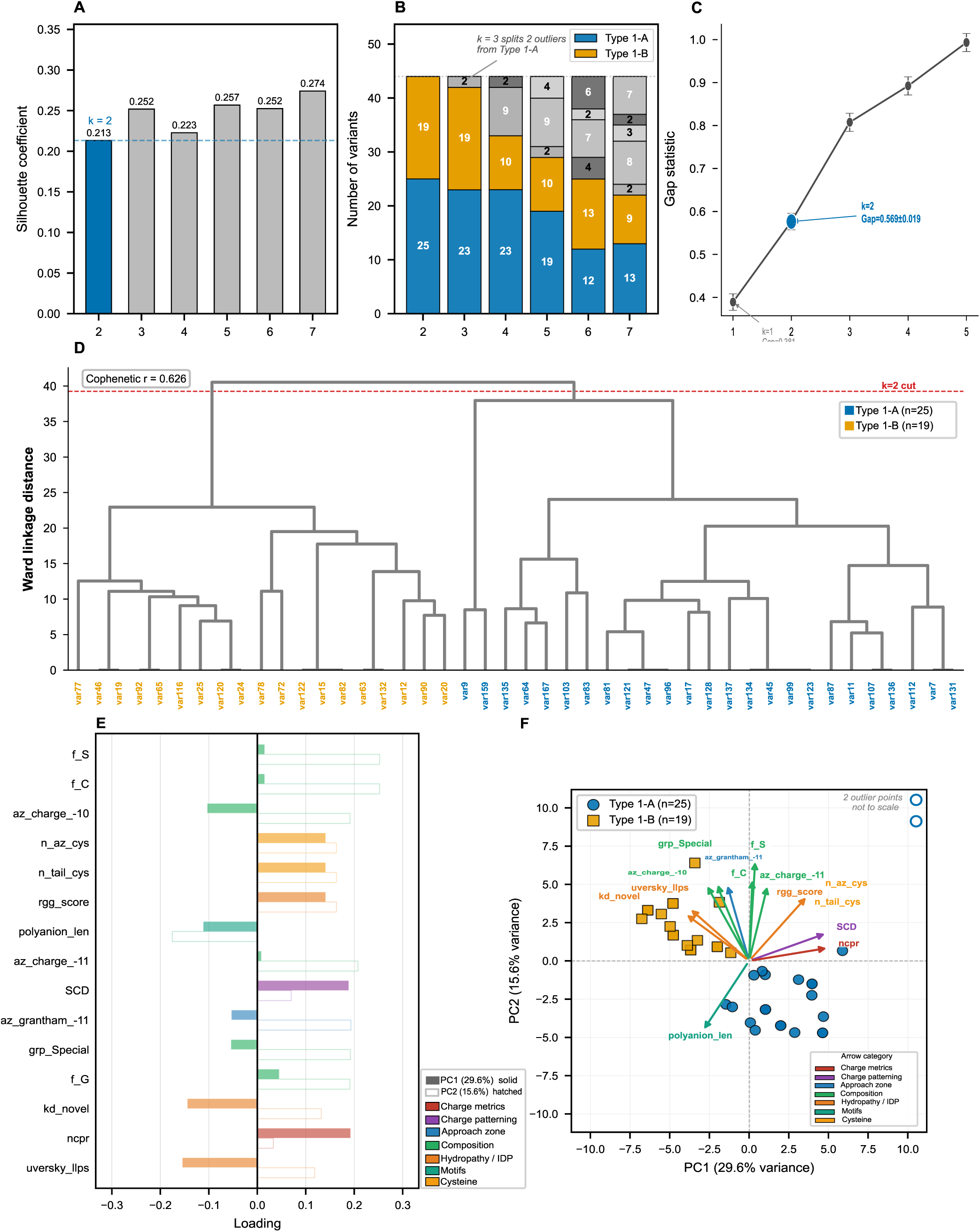
Clustering diagnostics and dimensionality reduction for unsupervised partitioning of Type 1-like variants (n = 44). (A) Silhouette coefficient for k = 2 to 7, computed by k-means clustering on 87 standardized sequence features. The k = 2 solution (silhouette = 0.213, highlighted in blue) was selected as the primary partition; values across all k range from 0.213 to 0.274. (B) Stacked bar chart showing cluster size composition at each k from 2 to 7, colored by Type 1-A (blue) and Type 1-B (orange) correspondence. At k = 2, clusters correspond to Type 1-A (n = 25) and Type 1-B (n = 19). At k = 3, two variants split from the Type 1-A cluster into a third group (n = 2), representing compositional outliers. The Type 1-B cluster remains stable from k = 3 onward; fragmentation occurs exclusively within the Type 1-A cluster at higher k. (C) Gap statistic for k = 1 to 5 (50 uniform reference datasets; error bars = SE). Gap at k = 1 (0.381 ± 0.019) is less than gap at k = 2 minus its SE (0.569 ± 0.019; threshold 0.550), satisfying the Tibshirani criterion and supporting k ≥ 2. (D) Ward hierarchical clustering dendrogram of all 44 Type 1-like variants. Leaf labels are variant identifiers; leaf colors indicate cluster membership (Type 1-A blue, Type 1-B orange). Red dashed line at Ward linkage distance ∼40 marks the cut yielding the two-cluster partition. Both major branches correspond cleanly to Type 1-A and Type 1-B; 100% membership agreement with k-means after label-permutation correction. Cophenetic correlation r = 0.626. (E) PCA loadings bar chart: top 15 sequence features by combined loading magnitude √(PC1² + PC2²). Solid bars = PC1 loadings (29.6% variance); hatched bars = PC2 loadings (15.6% variance). Bar colors indicate feature category: composition (green), hydropathy/IDP (orange), motifs (teal), charge patterning (purple), approach zone (blue), cysteine (yellow). (F) PCA biplot. Points colored by sequence subgroup (Type 1-A blue circles, Type 1-B orange squares). Loading arrows show the same top 15 features scaled to 60% of the display range, colored by category. Two outlier variants at PC1 > 10 shown as open symbols at clipped positions.

**Supplementary Figure S2.**
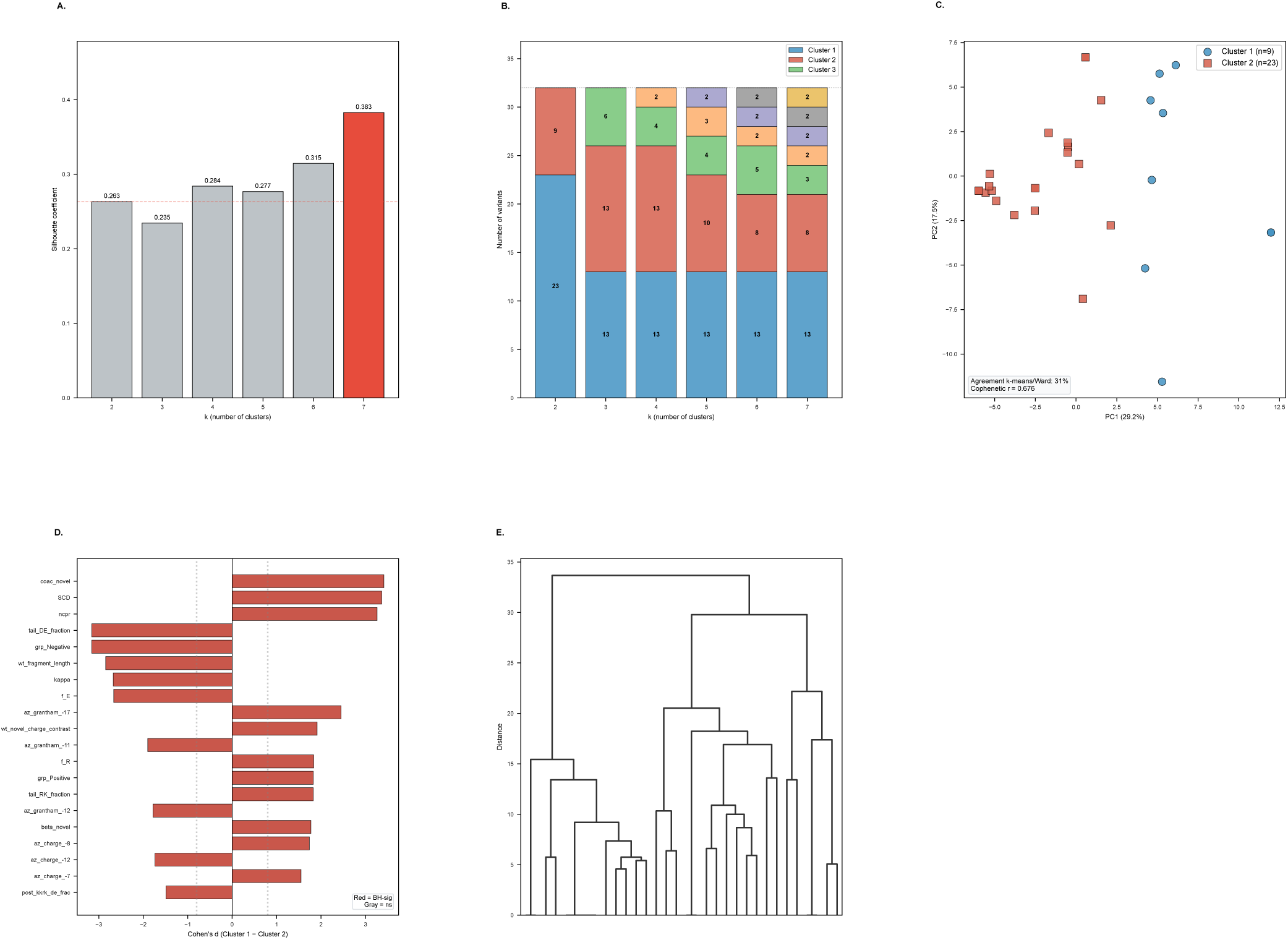
Unsupervised clustering analysis of Type 2 (ET) variants. All 32 Type 2 variants subjected to the same clustering pipeline as Type 1-like variants, using 85 standardized sequence features (KKRK_present excluded: zero variance in Type 2; Sγ–Sγ distance unavailable). (A) Silhouette coefficient k = 2 to 7. All values low (0.235–0.383); nominal best k = 7 (sil = 0.383) driven by progressive fragmentation into very small clusters. k = 2 silhouette = 0.263. (B) Cluster size composition at each k. At k = 2, the partition is 23 vs 9. From k = 3 onward, a stable cluster of 13 variants is consistently identified while remaining variants fragment into groups of 2–6, indicating a compositionally distinct minority rather than a cleanly separable second subgroup. (C) PCA of Type 2 variants colored by k = 2 cluster membership (PC1 = 29.2%, PC2 = 17.5%, combined 46.7%). K-means and Ward linkage agree on only 31% of assignments at k = 2, compared to 100% for Type 1-like variants, confirming absence of robust bipartite structure. Cophenetic r = 0.676. (D) Post-hoc feature importance for the k = 2 partition: 43 of 85 tested features BH-significant; top discriminators include NCPR, SCD, kappa, and Glu fraction, reflecting continuous charge variation rather than a discrete boundary. (E) Ward hierarchical dendrogram showing gradual, approximately equal-length branching without a dominant two-group partition, consistent with a largely homogeneous sequence architecture. Together, the low silhouette, poor k-means/Ward agreement, and dendrogram topology indicate that Type 2 variants do not harbor robust sequence-defined subgroups comparable to the Type 1-A/Type 1-B partition.

**Supplementary Figure S3.**
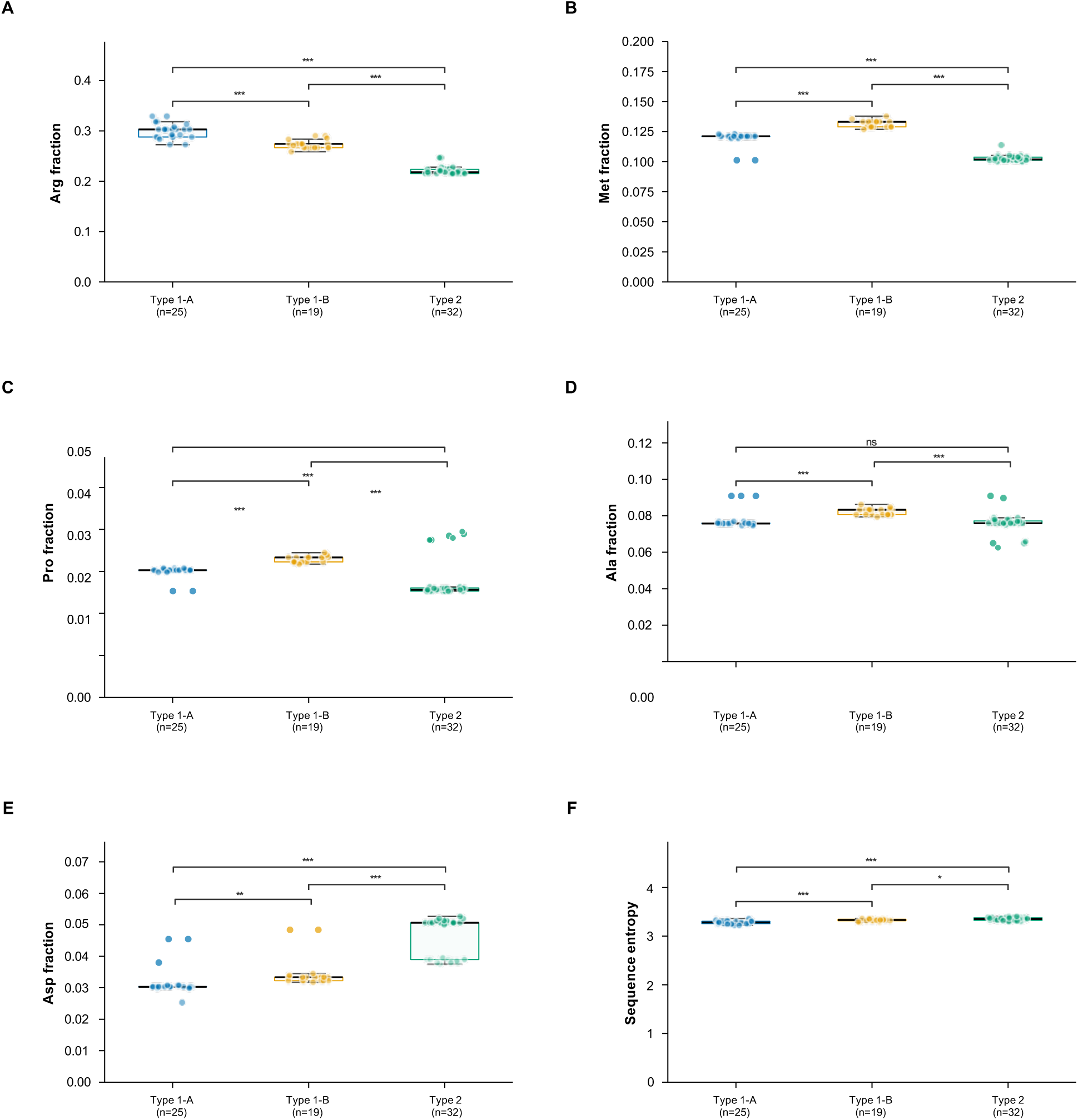
Amino acid composition of novel C-terminal tails across the three variant groups. (A) Arginine (Arg) fraction. Type 1-A highest (mean 0.300 ± 0.015), Type 1-B intermediate (0.273 ± 0.009), Type 2 lowest (0.221 ± 0.008); all pairwise ***p < 0.001. (B) Methionine (Met) fraction. Type 1-B highest (mean 0.132 ± 0.003), Type 1-A lower (0.120 ± 0.006), Type 2 lowest (0.103 ± 0.003); all pairwise ***p < 0.001. (C) Proline (Pro) fraction. Type 1-B highest (mean 0.033 ± 0.001), Type 1-A similar (0.030 ± 0.001), Type 2 lowest (0.028 ± 0.005); all pairwise ***p < 0.001. (D) Alanine (Ala) fraction. Type 1-A (mean 0.078 ± 0.005), Type 1-B (0.082 ± 0.002), Type 2 (0.076 ± 0.006); Type 1-A vs Type 2 ns; both comparisons with Type 1-B ***p < 0.001. (E) Aspartate (Asp) fraction. Type 2 substantially enriched (mean 0.046 ± 0.006) relative to Type 1-A (0.032 ± 0.005) and Type 1-B (0.035 ± 0.005); Type 1-A vs Type 1-B **p < 0.01; both vs Type 2 ***p < 0.001. (F) Sequence entropy (Shannon entropy, entropy_novel). Type 1-A lowest (mean 3.285 ± 0.033), Type 1-B intermediate (3.329 ± 0.025), Type 2 highest (3.350 ± 0.032); all pairwise comparisons significant. Box plots: median and IQR; whiskers 1.5× IQR; individual points shown. *p < 0.05; **p < 0.01; ***p < 0.001; ns, not significant (BH-corrected Mann-Whitney U).

**Supplementary Figure S4.**
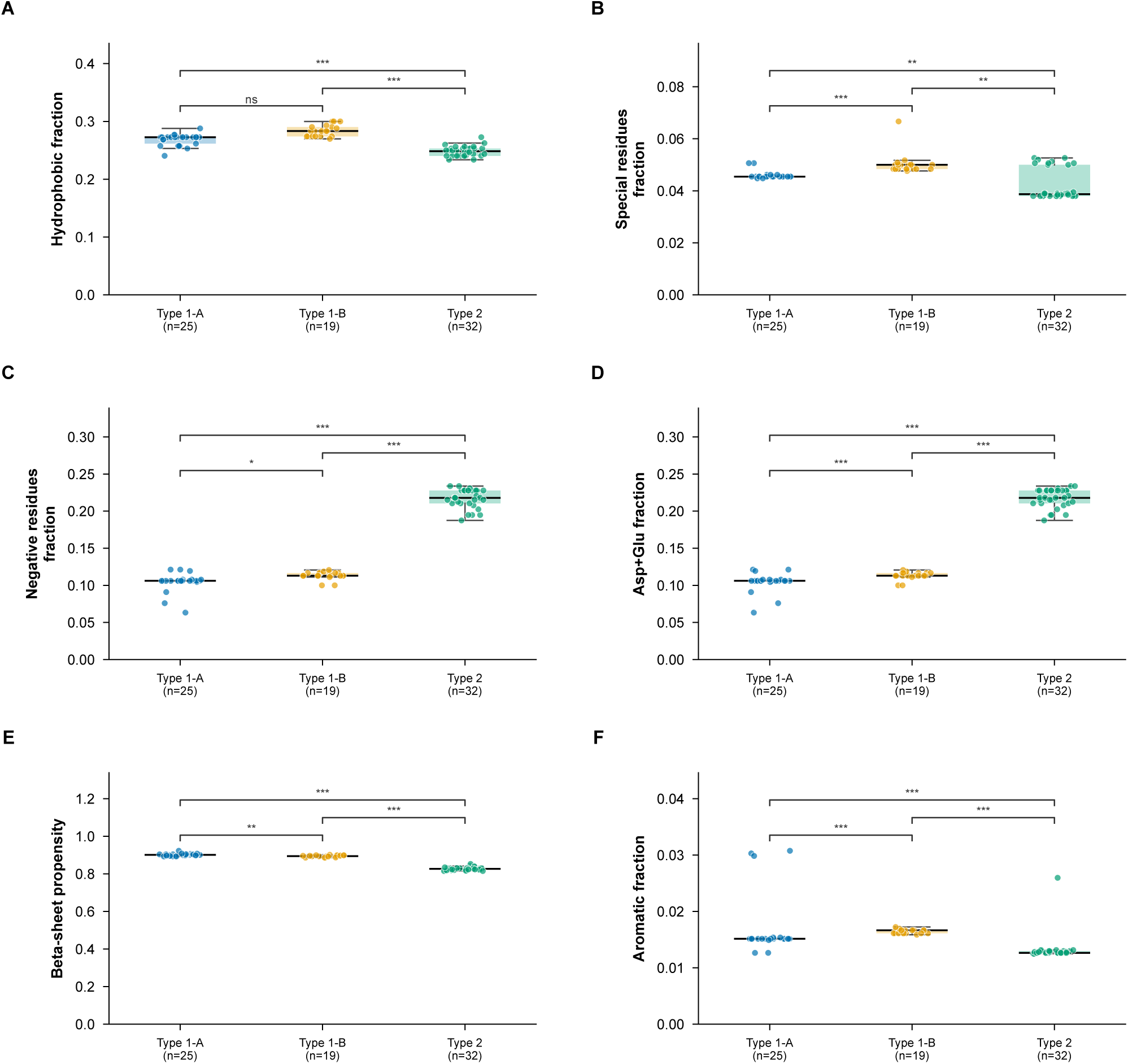
Residue group fractions and biophysical propensity scores across variant groups. (A) Hydrophobic residue fraction. Type 1-B highest (mean 0.284 ± 0.010), Type 1-A lower (0.268 ± 0.010), Type 2 lowest (0.248 ± 0.009); Type 1-A vs Type 1-B ns; both vs Type 2 ***p < 0.001. (B) Special residue fraction (Cys, Gly, Pro combined). Type 1-B slightly enriched (mean 0.050 ± 0.004) relative to Type 1-A (0.046 ± 0.002) and Type 2 (0.042 ± 0.006); Type 1-A vs Type 1-B ***p < 0.001; both vs Type 2 **p < 0.01. (C) Negative residue fraction (Asp + Glu). Type 2 dramatically enriched (mean 0.218 ± 0.013) relative to Type 1-A (0.104 ± 0.012) and Type 1-B (0.113 ± 0.005); Type 1-A vs Type 1-B *p < 0.05; both vs Type 2 ***p < 0.001. (D) Asp+Glu combined fraction. Type 1-A 0.104 ± 0.012, Type 1-B 0.113 ± 0.005, Type 2 0.218 ± 0.013; significance identical to panel C. (E) Chou-Fasman beta-sheet propensity. Type 1-A highest (mean 0.901 ± 0.007), Type 1-B similar (0.895 ± 0.005), Type 2 lowest (0.827 ± 0.008); Type 1-A vs Type 1-B **p < 0.01; both vs Type 2 ***p < 0.001. (F) Aromatic residue fraction. All groups low: Type 1-A 0.0168 ± 0.0051, Type 1-B 0.0164 ± 0.0004, Type 2 0.0132 ± 0.0023; Type 1-A vs Type 1-B ***p < 0.001; both vs Type 2 ***p < 0.001. Box plots: median and IQR; whiskers 1.5× IQR; individual points shown. *p < 0.05; **p < 0.01; ***p < 0.001; ns, not significant (BH-corrected Mann-Whitney U).

**Supplementary Figure S5.**
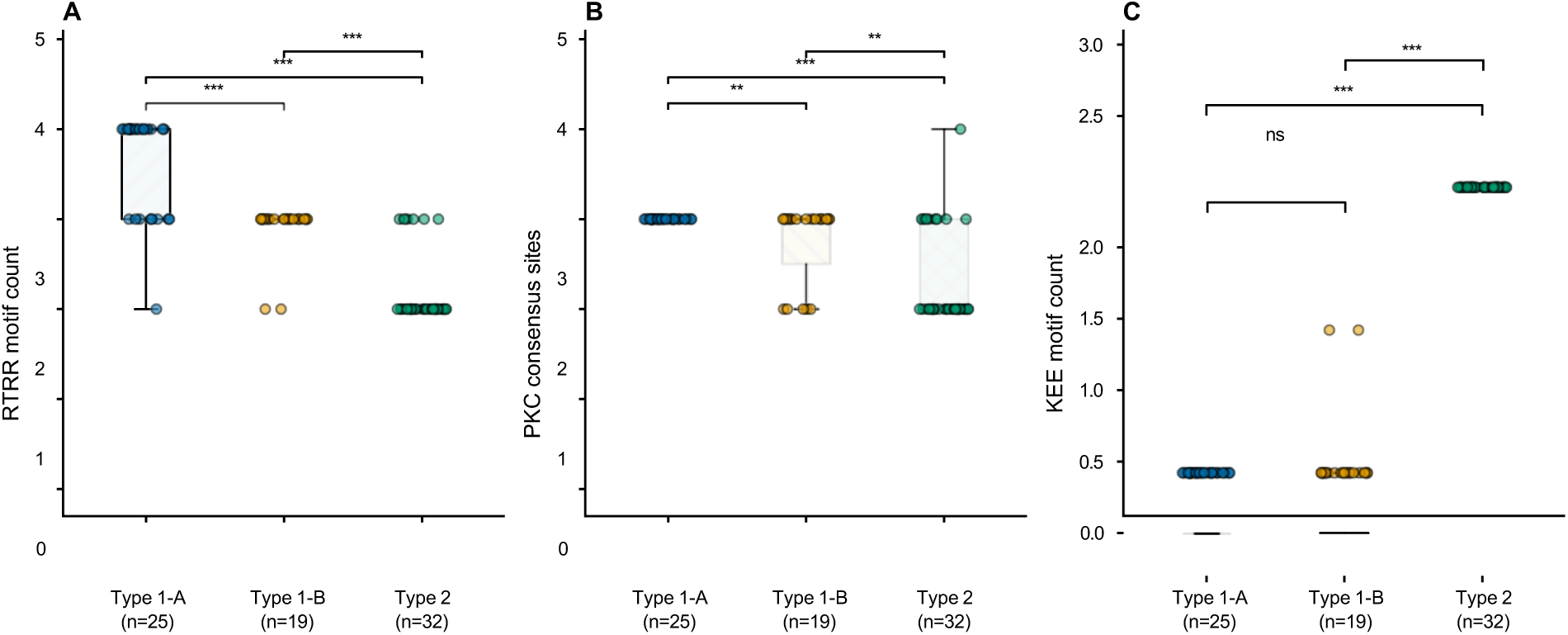
Sequence motif counts across variant groups. (A) RXRR motif count (R-any residue-R-R tetrapeptide) within the novel tail. Type 1-A highest (mean 3.600 ± 0.577, median 4), Type 1-B intermediate (2.895 ± 0.315, median 3), Type 2 lowest (2.188 ± 0.397, median 2); all pairwise ***p < 0.001. (B) Protein kinase C (PKC) consensus site count. Type 1-A invariant at 3 (mean 3.000 ± 0.000), Type 1-B intermediate (mean 2.737 ± 0.452, median 3), Type 2 lowest (mean 2.312 ± 0.535, median 2–3); Type 1-A vs Type 1-B **p < 0.01; both vs Type 2 ***p < 0.001. (C) KEE tripeptide motif count. Type 2 invariant at exactly 2 in all 32 variants (mean 2.000 ± 0.000); Type 1-A absent in all 25 variants (mean 0.000 ± 0.000); Type 1-B near-absent with 2 variants carrying KEE = 1 (mean 0.105 ± 0.315, median 0); Type 1-A vs Type 1-B ns; Type 1-A vs Type 2 ***p < 0.001; Type 1-B vs Type 2 ***p < 0.001. Box plots: median and IQR; whiskers 1.5× IQR; individual points shown. **p < 0.01; ***p < 0.001; ns, not significant (BH-corrected Mann-Whitney U).

**Supplementary Figure S6.**
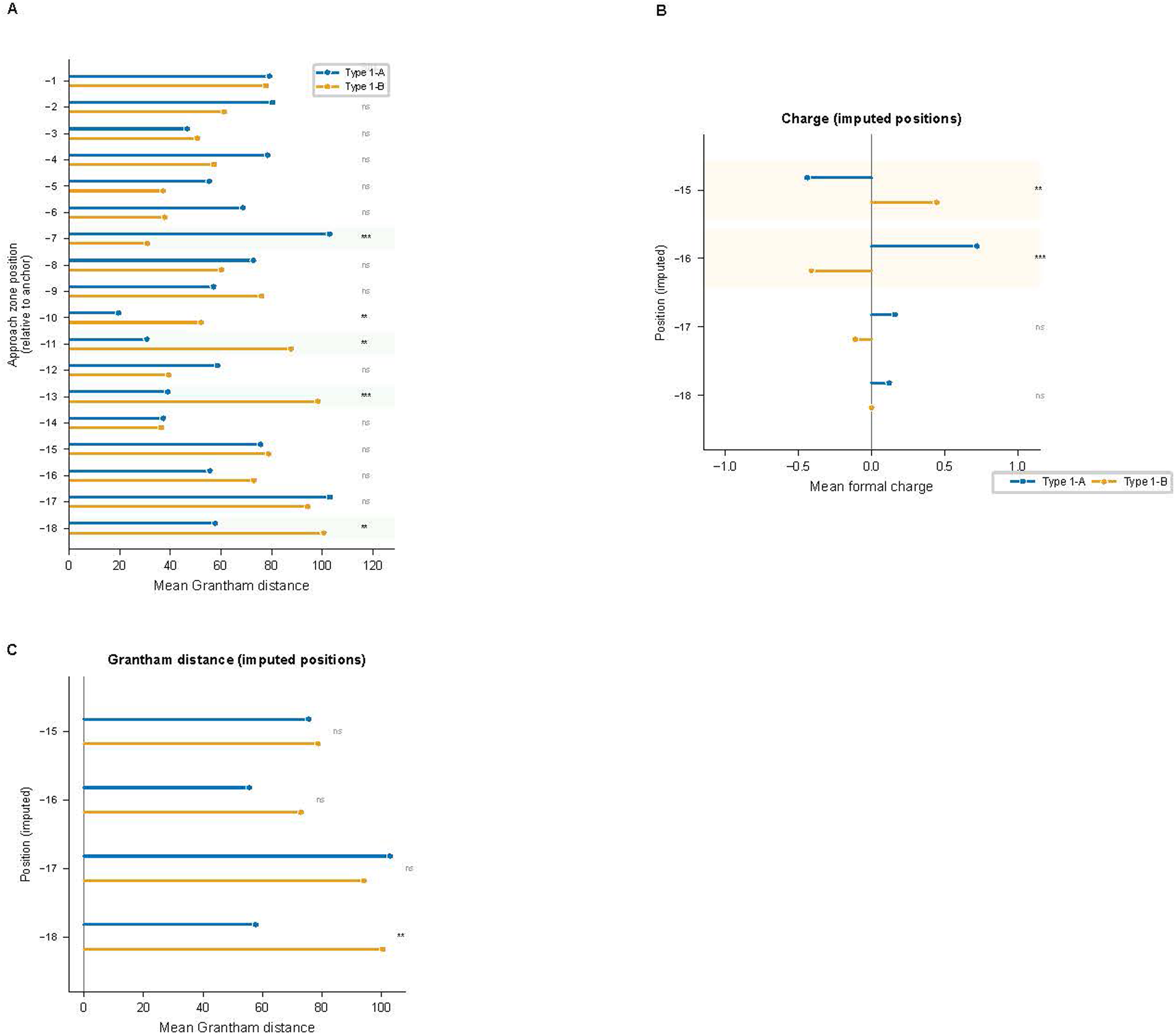
Approach zone physicochemical profiles across Type 1 subgroups. (A) Mean Grantham physicochemical distance at each approach zone position (positions −1 through −18, relative to anchor) for Type 1-A (blue) and Type 1-B (orange). Higher Grantham distance indicates greater physicochemical divergence from the population consensus residue at that position. Significant differences between subgroups are observed at positions −7 (***p < 0.001), −10 (**p < 0.01), −11 (**p < 0.01), −13 (***p < 0.001), and −18 (**p < 0.01); all remaining positions non-significant (BH-corrected Mann-Whitney U). (B) Mean formal charge at imputed approach zone positions −15 to −18 for Type 1-A (blue) and Type 1-B (orange). These positions are undefined for variants with shorter approach zones and were imputed with within-group feature medians prior to clustering (120 total imputed values, 3.1% of the data matrix). Positions −15 (**p < 0.01) and −16 (***p < 0.001) differ significantly; positions −17 and −18 are non-significant. (C) Mean Grantham distance at the same imputed positions −15 to −18. Position −18 is significant (**p < 0.01); positions −15 to −17 are non-significant. Values at imputed positions should be interpreted with caution. Positions −19 and −20 were excluded from all testing due to insufficient non-missing values. BH-corrected Mann-Whitney U.

**Supplementary Figure S7.**
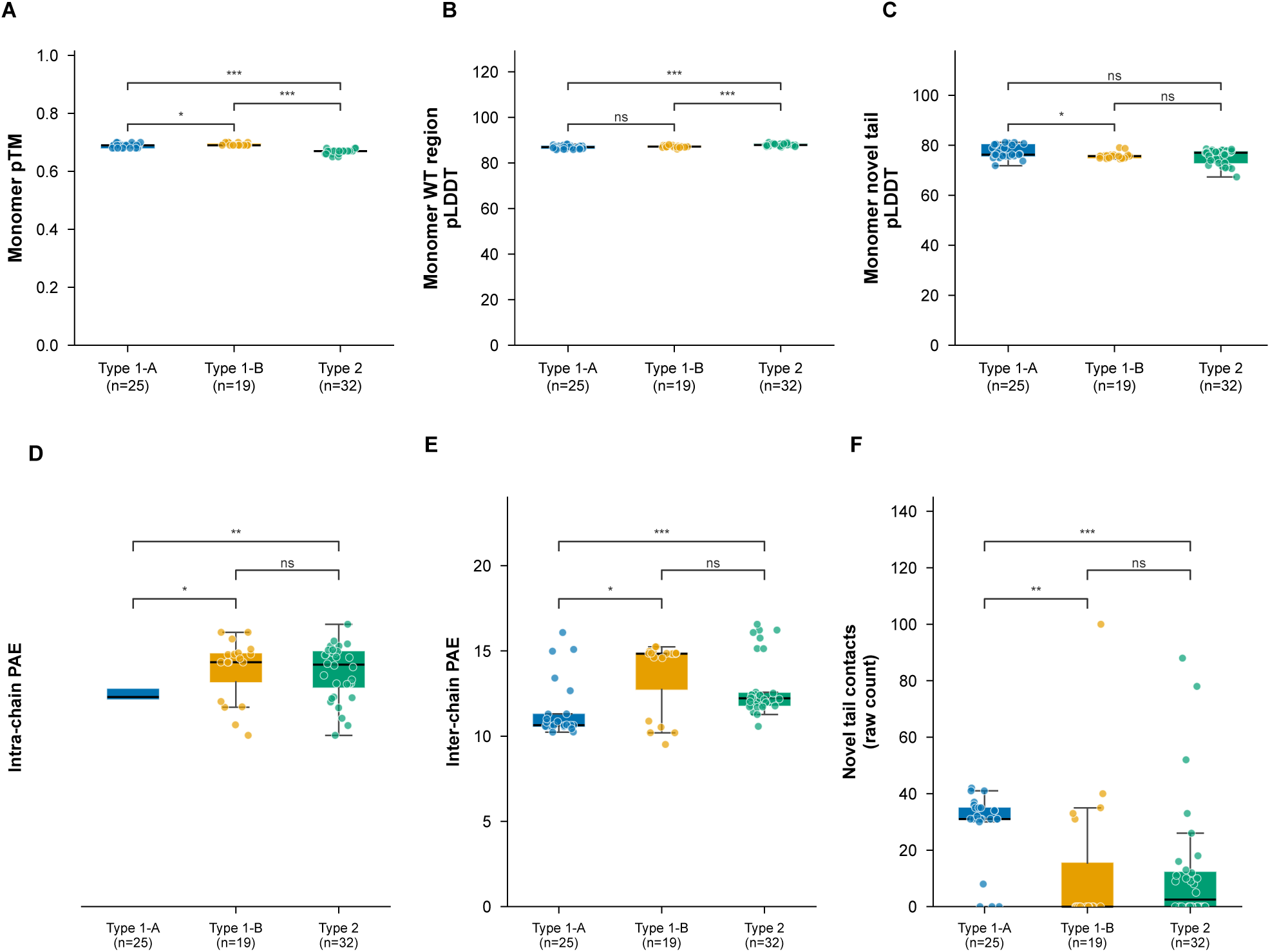
AlphaFold2 monomer and homodimer prediction confidence metrics. All metrics from unrelaxed rank-1 ColabFold v1.5 models. (A) Monomer predicted TM-score (mono_ptm). Type 1-A (mean 0.688 ± 0.007) and Type 1-B (0.693 ± 0.005) similar; Type 2 lower (0.668 ± 0.007); Type 1-A vs Type 1-B *p < 0.05; both vs Type 2 ***p < 0.001. (B) Monomer WT region pLDDT (mean pLDDT over residues upstream of frameshift). All groups high: Type 1-A 86.95 ± 0.75, Type 1-B 87.19 ± 0.44, Type 2 87.95 ± 0.52; Type 1-A vs Type 1-B ns; both vs Type 2 ***p < 0.001, confirming high confidence in the conserved domain across all variant groups. (C) Monomer novel tail pLDDT. Type 1-A highest (mean 77.43 ± 2.64), Type 1-B lower (75.79 ± 1.23), Type 2 similar to Type 1-B (75.46 ± 2.99); Type 1-A vs Type 1-B *p < 0.05; Type 1-B vs Type 2 ns; Type 1-A vs Type 2 ns. All values fall in the 70–90 range rather than the <50 band associated with intrinsically disordered regions in AlphaFold2, likely reflecting the conserved anchor providing a structured scaffold. (D) Intra-chain predicted aligned error (PAE; PAE between residues within the same chain). Type 1-A lowest (mean 11.43 ± 1.66), Type 2 intermediate (12.85 ± 1.69), Type 1-B highest (13.68 ± 2.11); Type 1-A vs Type 1-B *p < 0.05; Type 1-A vs Type 2 ***p < 0.001; Type 1-B vs Type 2 ns. (E) Inter-chain PAE (PAE between residues on opposing chains). Type 1-A lowest (mean 23.20 ± 1.94, highest predicted interface confidence), Type 1-B (24.39 ± 2.96) and Type 2 (24.76 ± 3.50) higher; Type 1-A vs Type 1-B *p < 0.05; Type 1-A vs Type 2 **p < 0.01; Type 1-B vs Type 2 ns. (F) Raw (unnormalized) inter-chain novel tail contact count (Cα–Cα < 8 Å). Type 1-A highest (mean 28.92 ± 12.53, median 31); Type 1-B median 0 (mean 12.58 ± 25.68); Type 2 median 2.5 (mean 12.44 ± 21.86); Type 1-A vs Type 1-B **p < 0.01; Type 1-A vs Type 2 ***p < 0.001; Type 1-B vs Type 2 ns. Box plots: median and IQR; whiskers 1.5× IQR; individual points shown. *p < 0.05; **p < 0.01; ***p < 0.001; ns, not significant (BH-corrected Mann-Whitney U).

**Supplementary Figure S8.**
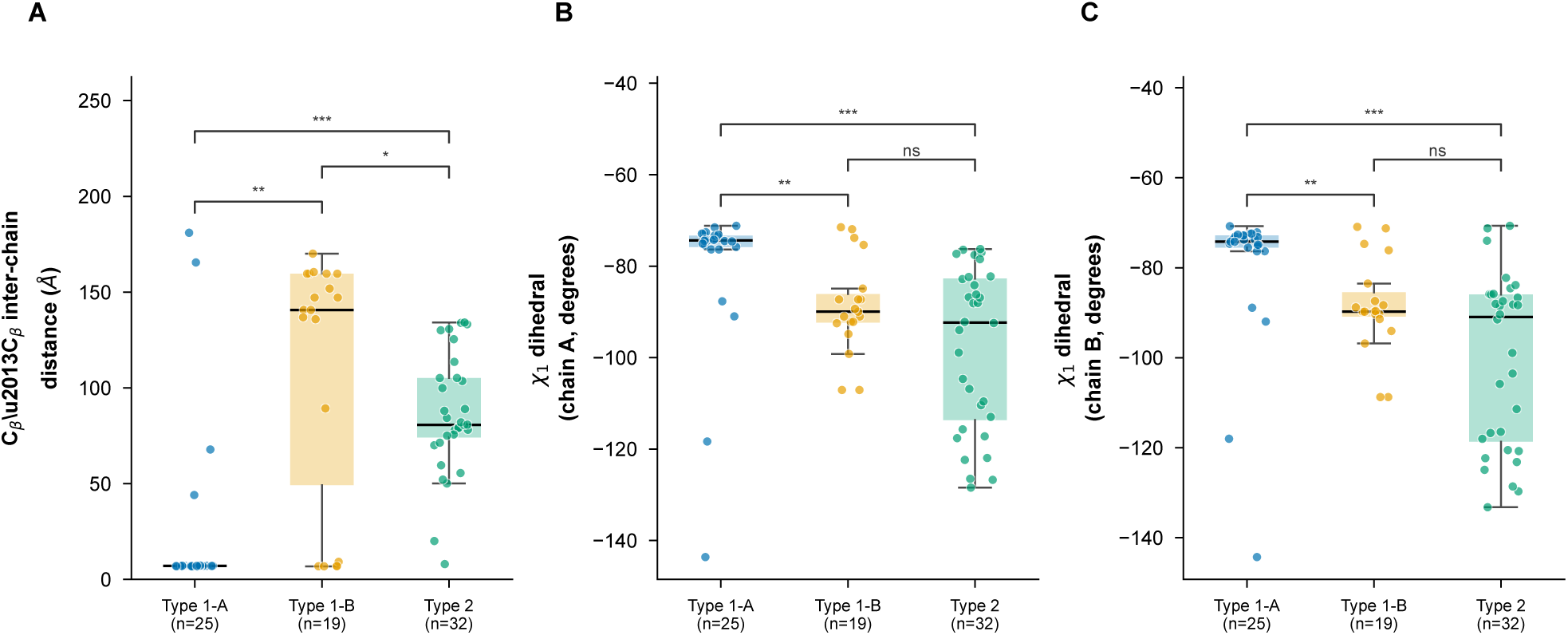
Cβ–Cβ inter-chain distance and χ1 dihedral angles at the anchor+32 cysteine. All metrics from unrelaxed rank-1 ColabFold v1.5 homodimer models at the conserved cysteine at anchor-relative position +32. (A) Cβ–Cβ inter-chain distance (Å). Type 1-A shows compact geometry (mean 24.18 ± 47.03 Å, median 7.02 Å), reflecting the bimodal distribution dominated by compact values; Type 1-B shows widely separated chains (mean 110.30 ± 65.36 Å, median 140.67 Å); Type 2 intermediate (mean 85.98 ± 31.05 Å, median 80.72 Å). Cβ–Cβ distance is highly correlated with Sγ–Sγ distance (Spearman r = 0.996). Type 1-A vs Type 1-B **p < 0.01; Type 1-A vs Type 2 ***p < 0.001; Type 1-B vs Type 2 *p < 0.05. (B) χ1 dihedral angle at the anchor+32 cysteine, Chain A. Type 1-A least negative (mean −79.80 ± 16.44°, median −74.36°); Type 1-B more negative (mean −88.72 ± 10.20°, median −89.95°); Type 2 most negative (mean −97.79 ± 17.61°, median −92.32°); Type 1-A vs Type 1-B **p < 0.01; Type 1-B vs Type 2 ns; Type 1-A vs Type 2 ***p < 0.001. (C) χ1 dihedral angle, Chain B. Pattern mirrors Chain A: Type 1-A mean −79.72 ± 16.65° (median −74.20°); Type 1-B mean −88.44 ± 10.32° (median −89.77°); Type 2 mean −100.23 ± 19.00° (median −90.93°); same significance pattern as (B). Box plots: median and IQR; whiskers 1.5× IQR; individual points shown. *p < 0.05; **p < 0.01; ***p < 0.001; ns, not significant (BH-corrected Mann-Whitney U).

**Supplementary Figure S9.**
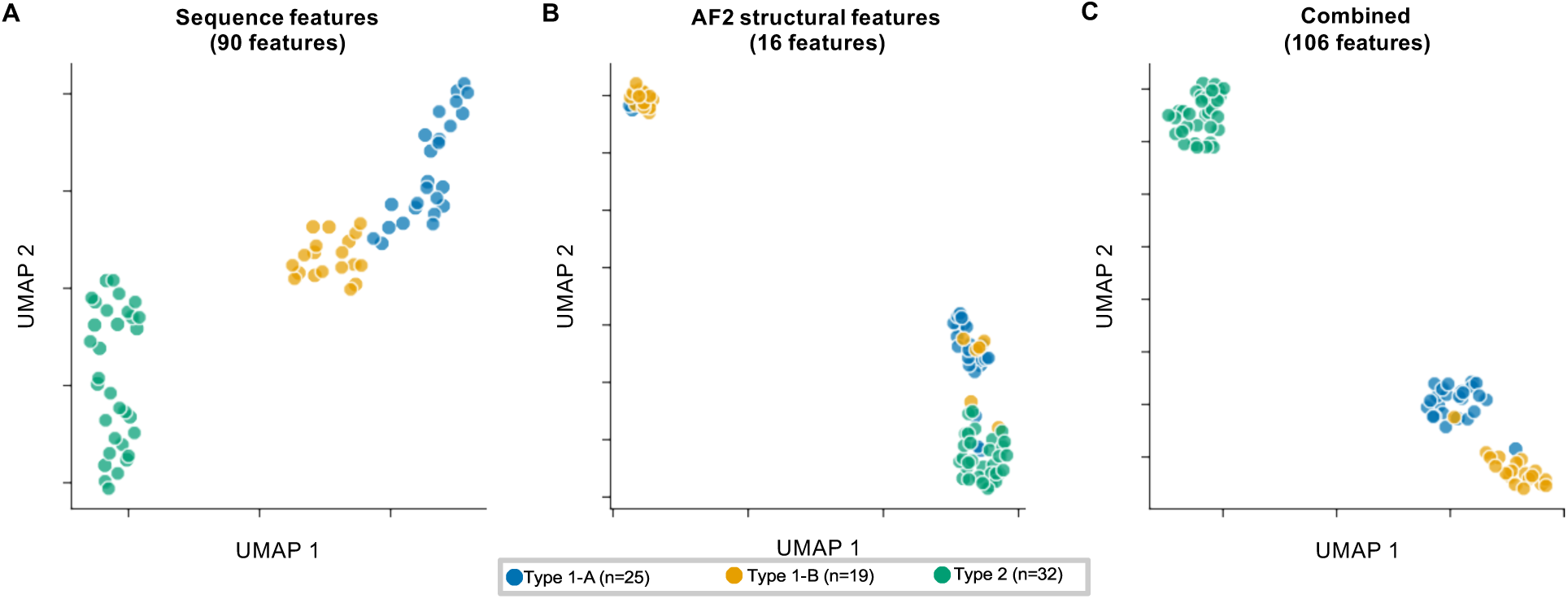
UMAP dimensionality reduction of all 76 variants across three feature spaces. UMAP embeddings computed independently for each feature space (n_neighbors = 12, min_dist = 0.3, metric = Euclidean, random_state = 42); all 76 variants shown in each panel, colored by variant group (Type 1-A blue, Type 1-B orange, Type 2 teal). (A) Sequence features (90 features). Type 1-A and Type 1-B form an overlapping continuum, well-separated from a compact Type 2 cluster. (B) AF2 structural features (16 features). Three groups form tighter, better-separated clusters; Type 1-A and Type 1-B partially overlap but are offset from each other, and both are clearly separated from Type 2. (C) Combined sequence and AF2 features (106 features). Three discrete clusters emerge with minimal overlap: Type 2 fully separated, Type 1-A and Type 1-B forming adjacent but distinct groups. Axes are in arbitrary UMAP units and are not directly comparable across panels.

**Supplementary Figure S10.**
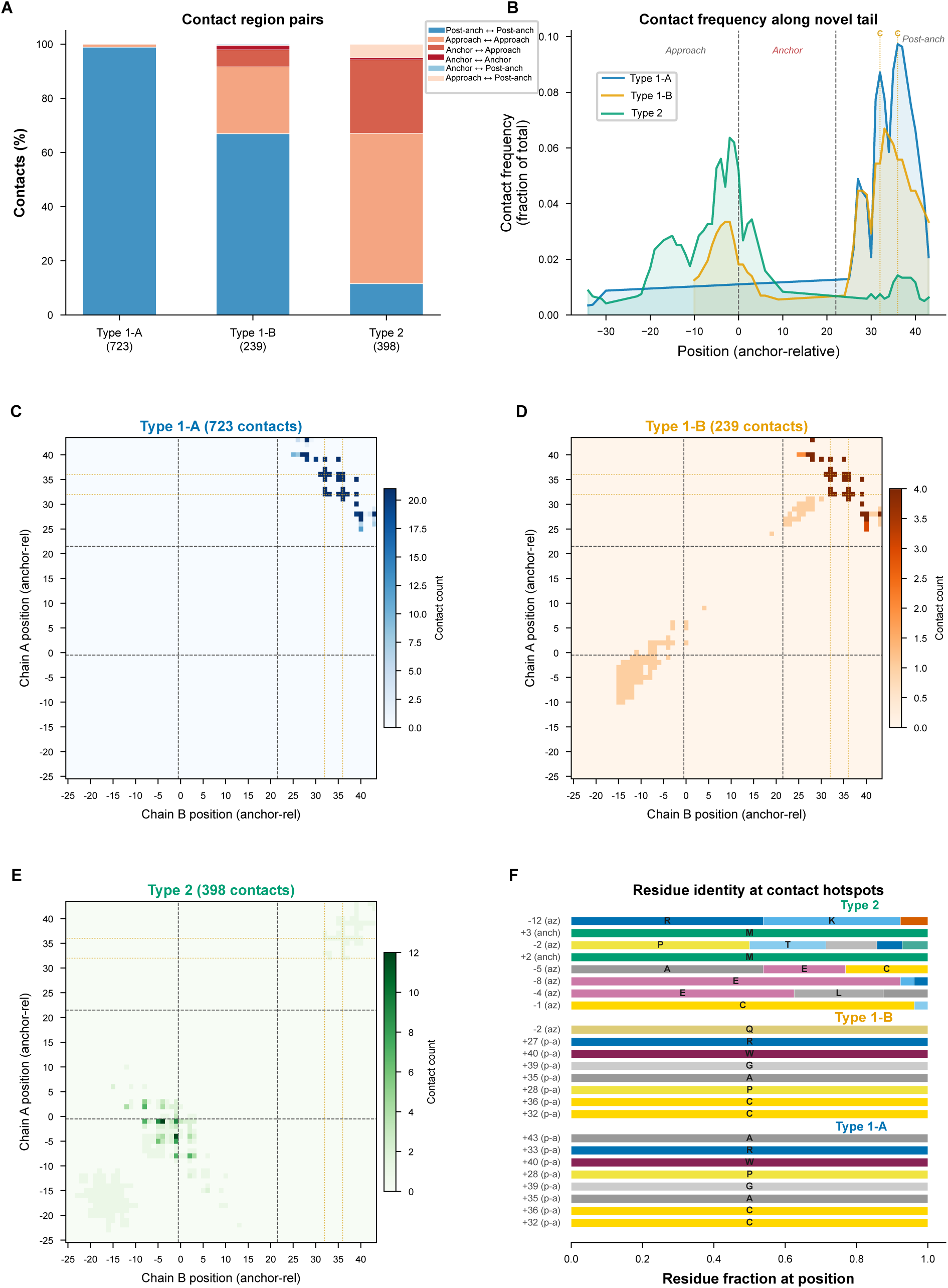
Inter-chain contact topology of predicted CALR homodimers. All inter-chain Cα–Cα contacts (< 8 Å cutoff) within the novel C-terminal tail extracted from unrelaxed rank-1 ColabFold v1.5 homodimer models for all 76 variants. Positions expressed relative to the anchor sequence (position 0 = first anchor residue; negative = approach zone; positive = post-anchor). (A) Contact region pair distribution for each group (total contacts: Type 1-A = 723, Type 1-B = 239, Type 2 = 398). Type 1-A contacts are 98.9% post-anchor ↔ post-anchor; Type 2 contacts are 55.5% approach ↔ approach and 27.1% anchor ↔ approach; Type 1-B shows a mixed pattern (66.9% post-anchor ↔ post-anchor, 24.7% approach ↔ approach). (B) Contact frequency profile (fraction of total contacts per position) along the novel tail for each group. Type 1-A shows a sharp peak at post-anchor positions +32 and +36 (invariant cysteines, together accounting for 32.1% of all Type 1-A contacts); Type 2 peaks in the approach zone near position −1; Type 1-B shows a low, mixed profile. Vertical dashed lines demarcate approach zone, anchor, and post-anchor regions; orange dashed lines mark cysteine positions +32 and +36. (C–E) Pairwise inter-chain contact maps for Type 1-A (C, blue, 723 contacts), Type 1-B (D, brown, 239 contacts), and Type 2 (E, green, 398 contacts). Each cell represents the total contact count between Chain A position (y-axis) and Chain B position (x-axis). Type 1-A contacts concentrate in the upper-right post-anchor quadrant; Type 1-B contacts are sparse with weak post-anchor signal; Type 2 contacts concentrate near the origin (approach zone). Color scales differ across panels (Type 1-A: 0–20; Type 1-B: 0–4; Type 2: 0–12). (F) Residue identity at the top contact hotspot positions for each group. Bar length indicates the fraction of variants with a given residue at that position. Type 1-A hotspots are dominated by the invariant cysteines at post-anchor positions +32 and +36. Type 2 hotspots are at approach zone positions −1, −4, −5, and −8; position −1 is the highest-contact position (53 contacts total), with cysteine the predominant residue in 59% of Type 2 variants, though contact frequency at this position does not correlate with cysteine identity. Type 1-B hotspots are at post-anchor positions +32 and +36 but with substantially lower contact counts than Type 1-A.

**Supplementary Figure S11.**
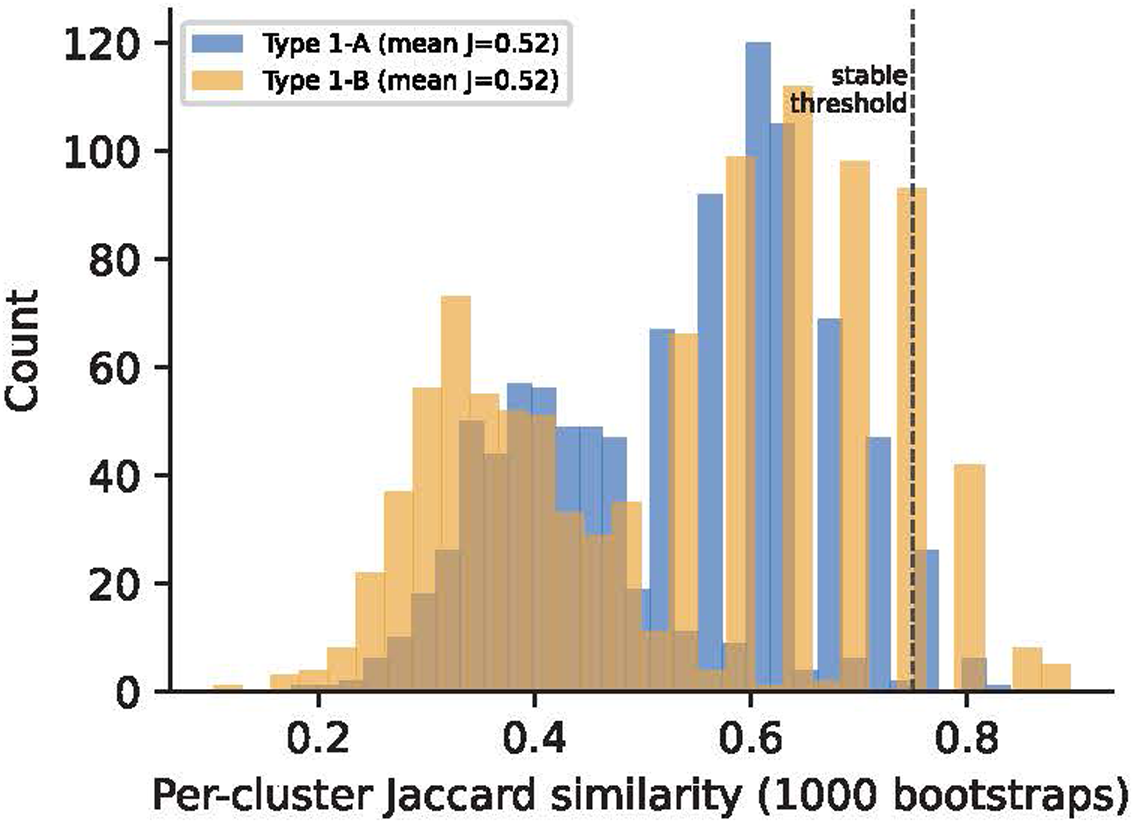
Bootstrap stability of the Type 1-A/Type 1-B partition. Distribution of per-cluster Jaccard similarity over 1,000 bootstrap resamples (Type 1-A blue, Type 1-B orange). Both clusters have mean Jaccard ≈ 0.52 (Type 1-A 0.523, Type 1-B 0.516), below the conventional stability threshold (0.75, dashed line; reached by 3% and 6% of resamples respectively), indicating the membership boundary is not robust to resampling and supporting interpretation of the subgroups as soft-bounded regions of a continuum rather than discrete clusters.

**Supplementary Figure S12.**
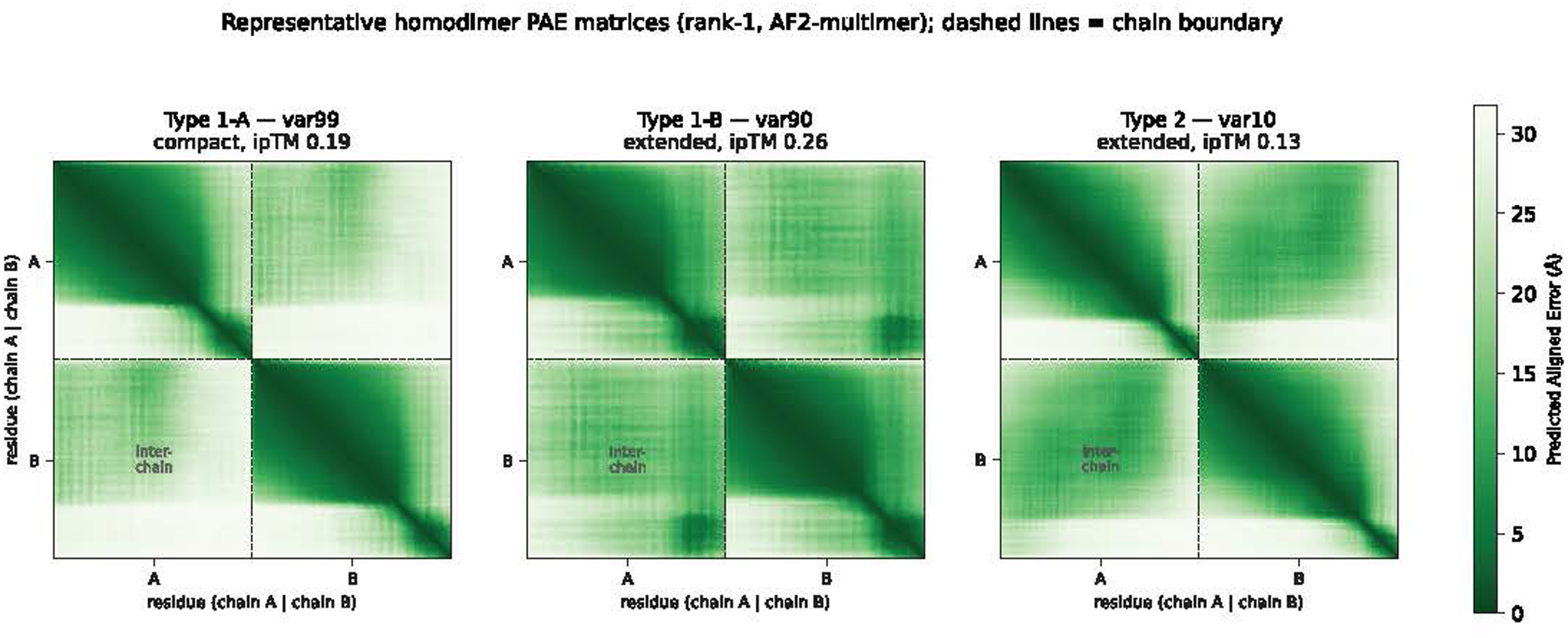
Representative homodimer PAE matrices. Predicted aligned error for one representative variant per group (Type 1-A var99, compact; Type 1-B var90, extended; Type 2 var10, extended; rank-1 models). Dashed lines mark the chain boundary. Inter-chain (off-diagonal) mean PAE (24.7/17.7/19.0 Å for the three variants) substantially exceeds intra-chain values (12.3/10.2/13.2 Å), indicating each chain is individually well-predicted while the interface is uncertain.

